# Role of c-Met/β1 integrin complex in the metastatic cascade

**DOI:** 10.1101/2020.04.20.051888

**Authors:** Darryl Lau, Harsh Wadhwa, Sweta Sudhir, Saket Jain, Ankush Chandra, Alan Nguyen, Jordan Spatz, Sumedh S. Shah, Justin Cheng, Michael Safaee, Garima Yagnik, Arman Jahangiri, Manish K. Aghi

## Abstract

Metastases cause 90% of human cancer deaths. The metastatic cascade involves local invasion, intravasation, extravasation, metastatic site colonization, and proliferation. While individual mediators of these processes have been investigated, interactions between these mediators remain less well defined. We previously identified a structural complex between receptor tyrosine kinase c-Met and β1 integrin in metastases. Using novel cell culture and *in vivo* assays, we found that c-Met/β1 complex induction promotes breast cancer intravasation and adhesion to the vessel wall, but does not increase extravasation. These effects may be driven by the ability of the c-Met/β1 complex to increase mesenchymal and stem cell characteristics. Multiplex transcriptomic analysis revealed upregulated Wnt and hedgehog pathways after c-Met/β1 complex induction. A β1 integrin point mutation that prevented binding to c-Met reduced intravasation. OS2966, a therapeutic antibody disrupting c-Met/β1 binding, decreased invasion and mesenchymal gene expression and morphology of breast cancer cells. Bone-seeking breast cancer cells exhibited higher c-Met/β1 complex levels than parental controls and preferentially adhere to tissue-specific matrix. Patient bone metastases demonstrated higher c-Met/β1 levels than brain metastases. Thus, the c-Met/β1 complex drives breast cancer cell intravasation and preferential affinity for bone tissue-specific matrix. Pharmacological targeting of the complex may prevent metastases, particularly osseous metastases.

## INTRODUCTION

Metastatic spread of cancer, one of the hallmarks of malignancy, is the main cause of up to 90% of human cancer deaths (1) and confers a median survival of less than six months once identified (2). Despite the clinical importance and tremendous public health impact of metastases, there remains insufficient understanding of the underlying molecular mechanisms of metastases and, importantly, this results in a lack of potential targetable vulnerabilities of the metastatic process.

The metastatic cascade comprises five major steps: local invasion of tumor cells at the primary site, transendothelial migration of tumor cells into the circulation (intravasation), egress out of the circulation and into sites of intended metastasis (extravasation), colonization at the metastatic site, and proliferation at the metastatic site leading to clinically detectable metastases (3). The role of specific mediators in early versus later steps of the metastatic cascade remains in need of clarification. The metastatic process is highly inefficient, as only 0.01% of cancer cells released into the circulation develop into metastatic foci (4), underscoring the importance of identifying the most vulnerable steps of the cascade and developing therapies targeting those steps.

There is evidence suggesting the importance of the latter steps in the cascade dating back to over a century ago, when Paget suggested that metastasis is not due to chance, but rather that some tumor cells (the “seed”) grow preferentially in the microenvironment of select organs (the “soil”) and that metastases result only from the appropriate seed in suitable soil (5). More modern gene expression studies have supported the hypothesis that tumor cells gain “metastasis virulence genes” that fail to affect primary tumor development and confer survival advantages only within the context of specific foreign microenvironments (6).

While individual mediators of these processes have been investigated, the interactions between these mediators have been less well studied. We previously demonstrated increased formation of a structural complex between receptor tyrosine kinase c-Met and β1 integrin in metastases compared to primary tumors (7). Here, we used novel cell culture models and *in vivo* assays to reveal the role of the c-Met/β1 complex in specific steps of the metastatic cascade in breast cancer.

The data presented in this study shows that the c-Met/β1 complex drives breast cancer cell intravasation and induces preferential affinity for tissue-specific matrix, particularly for osseous sites. This phenotype is secondary to changes is mesenchymal gene expression profile and increased stem cell characteristics. These findings reveal the c-Met/β1 complex to be a viable pharmacologic target to prevent metastases.

## RESULTS

### The c-Met/β1 complex promotes expression of genes related to cancer pathways and progression in breast cancer cells

We began by investigating the downstream effects of c-Met/β1 complex formation in breast cancer cells. To do so, we used our previously described MDA-MB-231-iDimerize-c-Met-β1 cells, which we engineered to express β1 integrin and c-Met fused to FRB (DmrC) and FKBP (DmrA), respectively, which enables us to regulate c-Met/β1 complex formation using AP21967 (A/C ligand heterodimerizer), a derivative of rapamycin (8). Gene expression changes induced by AP21967 treatment in MDA-MB-231-iDimerize-c-Met-β1 cells were then assessed in the NanoString nCounter platform using a 770 gene multiplex related to 13 cancer-associated canonical pathways (**Figures 1A-B; Supplemental Table 1; Supplemental Figure 1**). Pathway analysis revealed that AP21967 treatment upregulated genes in the Wnt and hedgehog pathways in MDA-MB-231-iDimerize-c-Met-β1 cells (**Figure 1C; Supplemental Figure 2; Supplemental Table 2**). AP21967 treatment also upregulated the expression of mesenchymal transcription factors TWIST (9), SNAIL (9), FOXC1 (10), FOXC2 (9), SLUG (11), and GSC (9) in MDA-MB-231-iDimerize-c-Met-β1 cells (P=0.013; **Figure 1D; Supplemental Figure 3**). We then performed a more downstream assessment in the NanoString nCounter platform by using a different multiplex to analyze expression of 770 genes from each step in the cancer progression process including angiogenesis, extracellular matrix (ECM) remodeling, epithelial-to-mesenchymal transition (EMT), and metastasis. This analysis revealed that AP21967 treatment upregulated the expression of genes in several downstream caner progression pathways in MDA-MB-231-iDimerize-c-Met-β1 cells (**Supplemental Figure 4**), including increased vasculogenesis, positive regulation of angiogenesis, hypoxia response, and stem cell associated scores (**Figure 1E**).

**Figure 1.**
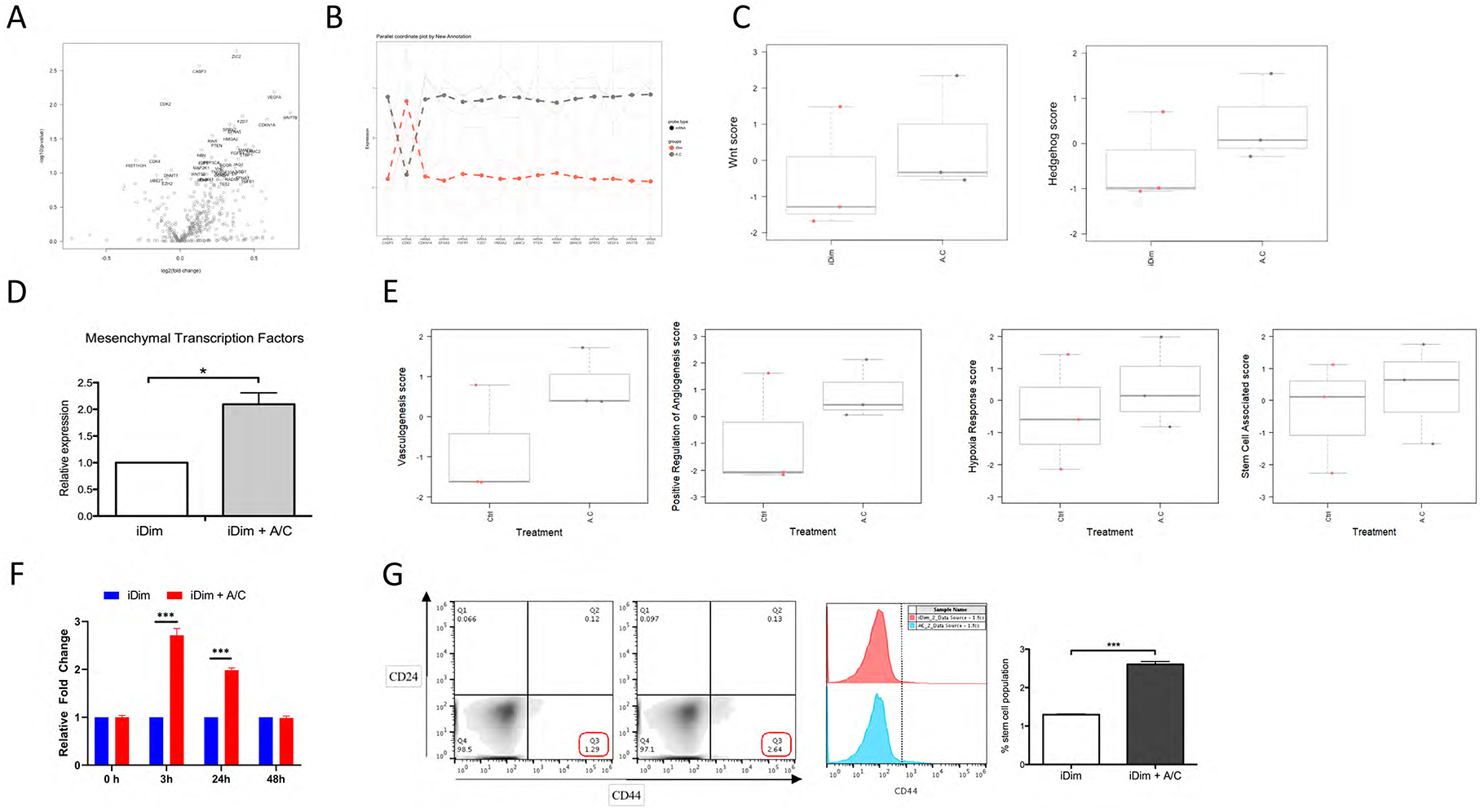
c-Met/β1 complex formation activates pathways implicated in metastases in breast cancer cells. Treatment of MDA-MB-231-iDimerize-c-Met-β1 cells with AP21967 induced c-Met/β1 complex formation, leading to upregulation of many of the 770 genes from 13 cancer-associated canonical pathways, as evidenced by (**A**) Volcano plot revealing the most upregulated genes with largest fold change in the upper right; (**B**) Top 15 hits based on p-values; and (**C**) Elevated expression of genes in the Wnt and Hedgehog pathways (n=3/group). (**D**) These pathways enriched the expression of mesenchymal transcription factors TWIST, SNAIL, FOXC1, FOXC2, SLUG, and GSC in AP21967-treated MDA-MB-231-iDimerize-c-Met-β1 cells, as assessed by qPCR (n=3/group; P=0.013). (**E**) Multiplex transcriptomic analysis revealed that AP21967 upregulated the expression of genes in several downstream caner progression pathways in MDA-MB-231-iDimerize-c-Met-β1, including increased vasculogenesis, positive regulation of angiogenesis, hypoxia response, and stem cell associated scores (n=3/group). AP21967 treatment also (**F**) increased the expression of stem cell genes c-Myc, Klf4, Oct4, Sox2, and Nanog (n=3/group; P=0.002) and (**G**) doubled the CD44+CD24-stem cell fraction of MDA-MB-231-iDimerize-c-Met-β1 cells (n=3/group; P<0.001). *P<0.05; **P<0.01; ***P<0.001.

### The c-Met/β1 complex enriches the stem cell fraction in breast cancer cells

Because of the findings of activated stem cell pathways from c-Met/β1 complex formation **(Figure 1E)**and because breast cancer stem cells have been shown to be a subset of breast cancer cells with enriched metastatic capacity (12), we then asked whether the ability of the c-Met/β1 complex to drive metastases (7) reflected an ability of the complex to enrich the breast cancer stem cell population. Indeed, AP21967 increased the expression of stem cell genes c-Myc (13), Klf4 (14), Oct4 (13), Sox2 (13), and Nanog (13) in MDA-MB-231-iDimerize-c-Met-β1 cells (P<0.001; **Figure 1F; Supplemental Figure 5**). Consistent with this gene expression profile, AP21967 also doubled the CD44+CD24-stem cell fraction (15) of MDA-MB-231-iDimerize-c-Met-β1 cells (P<0.001; **Figure 1G**; **Supplemental Figure 6**).

### The c-Met/β1 complex promotes intravasation of breast cancer cells

To determine the effects of the c-Met/β1 complex on intravasation of breast cancer cells, we developed a cell culture model of intravasation in which breast cancer cells were seeded in transwell chambers and allowed to sequentially traverse Matrigel and a HUVEC cell monolayer to model entrance into circulation (**Figure 2A**). This assay revealed that c-Met/β1 complex induction promoted intravasation of breast cancer cells (P=0.005; **Figure 2B**). Induction of the c-Met/β1 complex was also noted to increase adhesion of MDA-MB-231-iDimerize-c-Met-β1 cells to endothelial cells (P=0.004; **Figure 2C**). One potential mechanism of this c-Met/β1 complex-induced intravasation was noted when conditioned media from the breast cancer stem cells enriched by c-Met/β1 complex induction increased intravasation of breast cancer cells in our cell culture assay (P=0.0098; **Figure 2D; Supplemental Figure 7**). Because we previously showed that VEGF neutralizing antibody bevacizumab treatment increased c-Met/β1 complex formation in glioblastoma cells (7), we then analyzed the effects of bevacizumab on MDA-MB-231 cells. As with glioblastoma cells, bevacizumab increased c-Met/β1 complex formation in MDA-MB-231 cells (**Supplemental Figure 8**). Bevacizumab also increased expression of several of the pathway genes whose expression we showed to be driven by c-Met/β1 complex formation, including Wnt and hedgehog pathway gene Zic2 (P=0.003; **Figure 2E**). In confirmation of the functional consequences of bevacizumab-induced increased c-Met/β1 complex formation and increased cancer pathway gene expression, we found that bevacizumab increased intravasation of MDA-MB-231 cells (P<0.001; **Figure 2F; Supplemental Figure 9**).

**Figure 2.**
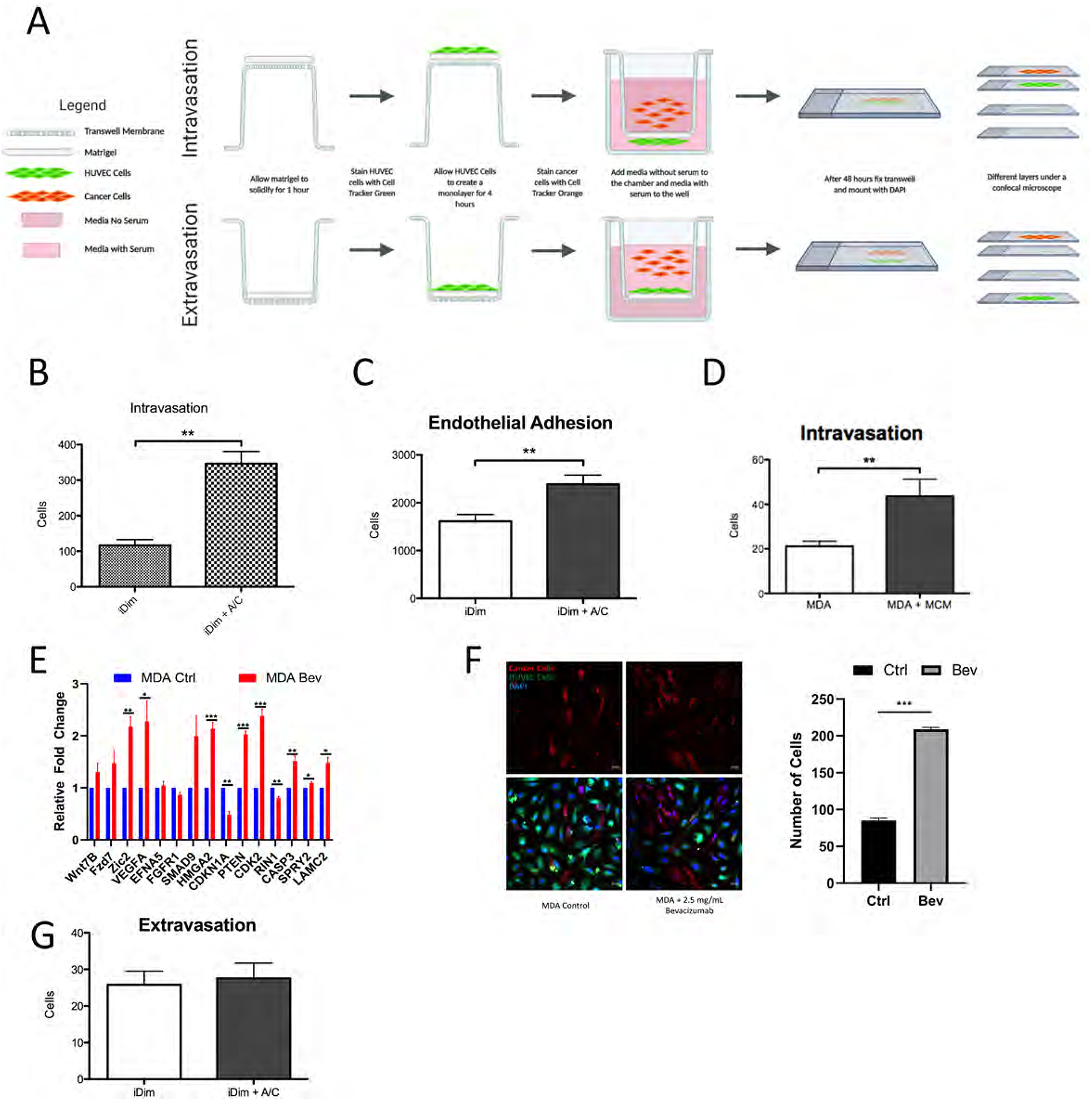
c-Met/β1 complex promotes intravasation of breast cancer cells. (**A**) Schema showing setup for cell culture assays that model breast cancer cell intravasation into and extravasation out of circulation. In these assays, Matrigel and a HUVEC monolayer are plated in orientation that allows modeling of intravasation of cancer cells into circulation and extravasation of cancer cells out of circulation. Induction of c-Met/β1 complex formation in MDA-MB-231-iDimerize-c-Met-β1 cells with AP21967 treatment increased (**B**) intravasation in the cell culture intravasation assay (n=3/group; P=0.005) and (**C**) adhesion of breast cancer cells to endothelial cells in cell culture (n=9/group; P=0.004). (**D**) Mammosphere conditioned media (MCM) increased intravasation of MDA-MB-231 breast cancer cells (n=9/group; P=0.0098). Bevacizumab increased (**E**) expression of several cancer signaling pathway genes, including Wnt and hedgehog pathway gene Zic2 (n=3/group; P=0.003) and (**F**) intravasation of MDA-MB-231 breast cancer cells (n=3/group; P<0.001). (**G**) Induction of c-Met/β1 complex formation in MDA-MB-231-iDimerize-c-Met-β1 cells with AP21967 treatment did not affect extravasation in the cell culture extravasation assay (n=9/group; P=0.8). *P<0.05; **P<0.01; ***P<0.001.

### The c-Met/β1 complex does not promote extravasation of breast cancer cells

To determine the effect of the c-Met/β1 complex on extravasation of breast cancer cells out of circulation, we modified our cell culture model of intravasation to make it model extravasation by seeding breast cancer cells in transwell chambers above a HUVEC cell monolayer and Matrigel (**Figure 2A**). This assay revealed that c-Met/β1 complex induction did not promote extravasation of breast cancer cells (P=0.8; **Figure 2G; Supplemental Figure 10**).

### The c-Met/β1 complex promotes tissue-specific metastases of breast cancer cells

We then investigated whether the c-Met/β1 complex promotes tissue-specific metastases. We began by determining if the c-Met/β1 complex promoted adhesion to specific types of ECM proteins, and found that c-Met/β1 complex induction by AP21967 increased adhesion of MDA-MB-231-iDimerize-c-Met-β1 cells to collagen, fibronectin, or laminin (P<0.001; **Figure 3A**). We then assessed the effect of the c-Met/β1 complex on adhesion to purified collagen types I-IV and found that c-Met/β1 complex induction specifically increased adhesion of MDA-MB-231-iDimerize-c-Met-β1 cells to collagen type I (P=0.004) but not collagen types II, III, or IV (P=0.2-0.7; **Figure 3B**). We then sought to determine if c-Met/β1 complex levels varied in breast cancer cells with metastatic preference for specific organs depending on the collagen content of the target organ. To do so, we used proximity ligation assays to analyze levels of the c-Met/β1 complex in cells derived from MDA-MB-231 through serial *in vivo* metastases conferring affinity for specific organs (16–18). MDA-MB-231-BO bone-seeking cells had the greatest levels of complex, more than MDA-MB-231-LM2 lung-(P=0.04; **Figure 3C**) or MDA-MB-231-BR brain-(P<0.001; **Figure 3C**) seeking cells, while MDA-MB-231-LM2 lung-seeking cells also had more complex than MDA-MB-231-BR brain-seeking cells (P=0.04; **Figure 3C**). These elevated levels of complex in MDA-MB-231-BO bone-seeking cells compared to MDA-MB-231-LM2 lung-or MDA-MB-231-BR brain-seeking cells also led to greater levels of the Wnt/hedgehog signaling genes we identified as associated with the complex, with increased Wnt7B in MDA-MB-231-BO relative to MDA-MB-231-LM2 (P=0.01) or MDA-MB-231-BR (P=0.03); increased Zic2 in MDA-MB-231-BO relative to MDA-MB-231-BR (P=0.02); and increased Fzd7 in MDA-MB-231-BO relative to MDA-MB-231-LM2 (P<0.001) or MDA-MB-231-BR (P<0.001) (**Figure 3D**). These findings are consistent with the differential collagen affinities we identified since collagen type I is the predominant collagen type in bone (19) and lung (20), with greater levels in the former than the latter, while collagen type IV is the predominant collagen type in brain.

**Figure 3.**
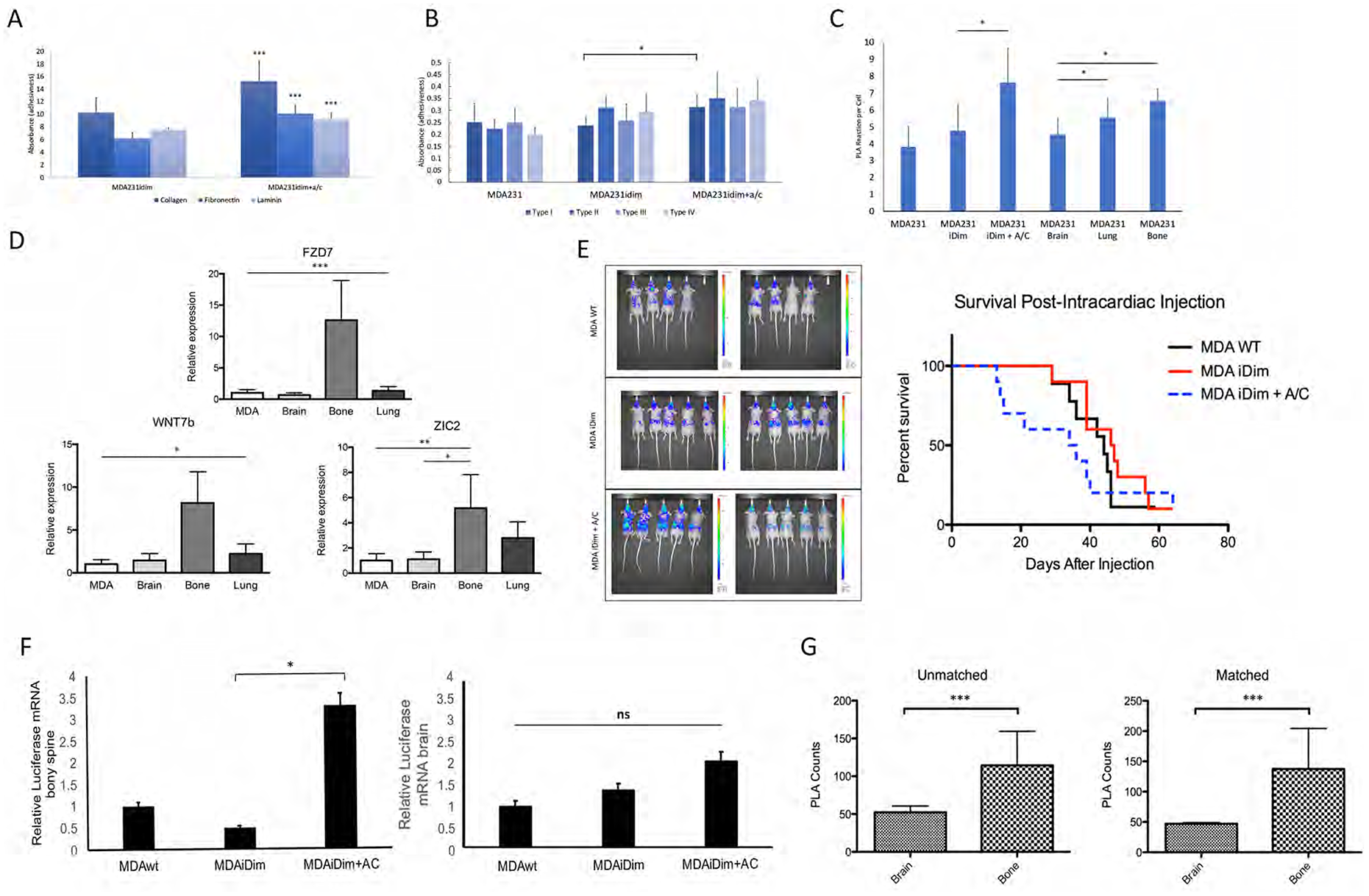
c-Met/β1 complex promotes affinity for specific ECM proteins in culture and tissue-specific metastases*in vivo*. **(A)**Induction of c-Met/β1 complex formation in MDA-MB-231-iDimerize-c-Met-β1 cells promoted adhesion to collagen, fibronectin, and laminin (n=32/group; P<0.001). (**B**) Induction of c-Met/β1 complex formation promoted adhesion of MDA-MB-231-iDimerize-c-Met-β1 cells to purified collagen type I (P=0.004) but not purified collagen types II, III, or IV (P=0.2-0.7) (n=32/group). (**C**) Proximity ligation assays revealed that cultured MDA-MB-231-BO bone-seeking cells had the greatest levels of c-Met/β1 complex, more than MDA-MB-231-LM2 lung-(P=0.04) or MDA-MB-231-BR brain-(P<0.001) seeking cells (n=9/group). (**D**) These greater levels of c-Met/β1 complex in MDA-MB-231-BO bone-seeking cells led to increased Wnt/hedgehog pathway signaling gene expression in these cells: increased Wnt7B expression in MDA-MB-231-BO relative to MDA-MB-231-LM2 (P=0.01) or MDA-MB-231-BR (P=0.03); increased Zic2 expression in MDA-MB-231-BO relative to MDA-MB-231-BR (P=0.02); and increased Fzd7 expression in MDA-MB-231-BO relative to MDA-MB-231-LM2 (P<0.001) or MDA-MB-231-BR (P<0.001) (n=3/group). (**E**) Luciferase-expressing MDA-MB-231-iDimerize-c-Met-β1 cells were pre-treated in culture with AP21967 versus vehicle, then implanted into athymic mice hearts, followed by serial treatment of mice with AP21967 or vehicle for the duration of mice survival while monitoring for metastases by bioluminescence. Shown to the left are bioluminescence imaging from the last time point before the first mouse died of tumor burden, and to the right are Kaplan-Meier curves from the 3 groups. We found that complex induction via AP21967 treatment in mice with MDA-MB-231-iDimerize-c-Met-β1 tumors shortened survival compared to vehicle treatment (n=8-10/group; P<0.001). (**F**) Micrometastases in specific end organs were detected by qPCR for the luciferase gene, with greater levels noted in the bony spine with AP21967 treatment of mice receiving intracardiac implantation of MDA-MB-231-iDimerize-c-Met-β1 tumor cells compared to without AP21967 treatment (n=3/group; P=0.02), but no significant changes noted in the brain (P=0.6). (**G**) PLA of patient metastases revealed increased c-Met/β1 complex in patient bony metastases (n=11) relative to brain metastases (n=12) from different patients (left panel; P<0.001) and in paired bone metastases relative to brain metastases from the same patients (n=3; right panel; P<0.001). *P<0.05; **P<0.01; ***P<0.001.

We then investigated the effects of the c-Met/β1 complex on tissue-specific metastases *in vivo* by pre-treating luciferase-expressing MDA-MB-231-iDimerize-c-Met-β1 cells in culture with AP21967 or vehicle, then performing intracardiac implantation of cells, followed by serial treatment of mice with AP21967 or vehicle for the duration of mice survival while monitoring for metastases by bioluminescence. We found that c-Met/β1 complex induction via AP21967 treatment resulted in significantly shorter survival (P<0.001; **Figure 3E**). While there was no difference in gross metastases detected by bioluminescence of AP21967 treated mice, more micrometastases to the bony spine (P=0.02), but not the brain (P=0.6), were noted with AP21967 treatment based on qPCR of these tissues for the luciferase gene expressed by the tumor cells (**Figure 3F**). Consistent with this *in vivo* finding and our finding that c-Met/β1 complex induction promoted adhesion specifically to collagen type I, levels of the complex assessed by proximity ligation assays (PLAs) were higher in patient metastases to the bony spine compared to patient metastases to the brain (P<0.001; **Figure 3G; Supplemental Figure 11**), with immunoprecipitation confirming this finding as well (**Supplemental Figure 12**).

### Genetically disrupting c-Met/β1 complex formation reduces the metastatic phenotype in breast cancer cells

Through PyMOL modeling and site-directed mutagenesis, we previously demonstrated that five individual amino acids in β1 integrin (246, 283, 284, 287, and 290) are crucial for binding of β1 integrin to c-Met (7). To determine the consequences of the lack of β1 integrin binding to c-Met, we utilized CRISPRi to knockout β1 integrin in MDA-MB-231 cells (**Supplemental Figure 13**) and then, via lentiviral transduction, restored wild type β1 integrin or β1 integrin with change of amino acid 246 or 287 from aspartate to alanine. Either of these point mutations reduced binding of β1 integrin to c-Met (**Supplemental Figure 14**) and β1D246A was chosen for further investigation. Multiplex expression analysis of 770 genes from each step in the cancer progression process including angiogenesis, ECM remodeling, epithelial-to-mesenchymal transition (EMT), and metastasis revealed that, compared to cells with restored wild-type β1 integrin, cells with β1D246A exhibited reduced expression of genes associated with cell motility and ECM structure and increased expression of genes associated with suppression of metastases and angiogenesis (**Figures 4A-C; Supplemental Table 3**). Functional changes corresponding to these alterations in gene expression were seen when, compared with MDA-MB-231 cells with restored wild-type β1 integrin, cells with β1D246A exhibited reduced intravasation (P<0.001; **Figure 4D**).

**Figure 4.**
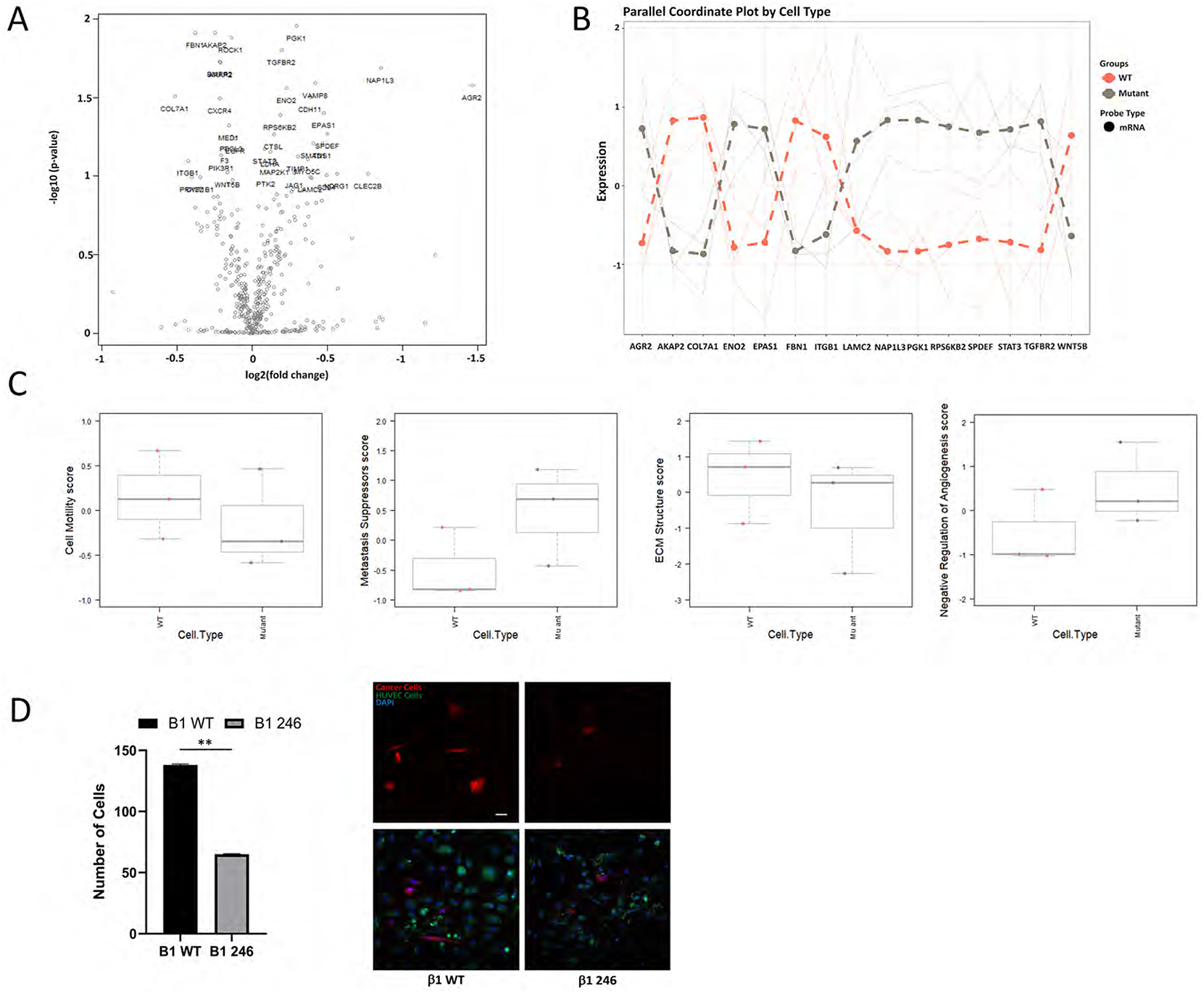
Genetic elimination of c-Met/β1 binding inhibits oncologic transcriptional changes and intravasation. Site-directed mutagenesis of β1 integrin to prevent binding to c-Met (the β1D246A mutation) in MDA-MB-231 cells altered expression of several of the 770 cancer progression-related genes in a multiplex panel, as evidenced by (**A**) Volcano plot revealing the most upregulated genes with largest fold change in the upper right; (**B**) Top 15 hits based on p-values; and (**C**) Box and whisker plots showing alterations in gene family scores. (**D**) These gene expression changes reduced intravasation in cell culture assays of MDA-MB-231 cells with β1D246A compared to MDA-MB-231 cells with wild-type β1 integrin (n=3/group; P=0.003). *P<0.05; **P<0.01; ***P<0.001.

### Pharmacologically targeting the c-Met/β1 complex reduces the metastatic phenotype in breast cancer cells

To target the c-Met/β1 complex in breast cancer cells, we treated cells with OS2966, a therapeutic humanized β1 integrin neutralizing antibody that we showed to inhibit c-Met/β1 complex formation in MDA-MB-231 cells (**Figure 5A**). AP21967 increased mesenchymal transcription factor expression in cultured MDA-MB-231-iDimerize-c-Met-β1 cells (P<0.001), changes reversed by OS2966 (P<0.001; **Figure 5B; Supplemental Figure 15**). Morphologic phenotypic effects associated with these transcription factor changes were also noted, as AP21967-induced c-Met/β1 complex formation lowered the form factor of MDA-MB-231-iDimerize-c-Met-β1 cells in culture (P=0.002, **Figure 5C**), consistent with a more mesenchymal morphology, and this effect was reversed by OS2966 (P<0.001; **Figure 5C**). Similarly, MDA-MB-231-BO and MDA-MB-231-BR organ-seeking cells had higher mesenchymal transcription factor expression than parental MDA-MB-231 cells, differences that were reversed with OS2966 treatment (P=0.008-0.009; **Figure 5D**). These organ-seeking cells also had lower form factor than parental MDA-MB-231 cells (P<0.001; **Figure 5E**), differences that were reversed with OS2966 treatment (P=0.007-0.03; **Figure 5E**). OS2966 also reversed the invasiveness (P=0.04; **Figure 5F**) caused by AP21967 treatment of MDA-MB-231-iDimerize-c-Met-β1 cells in culture.

**Figure 5.**
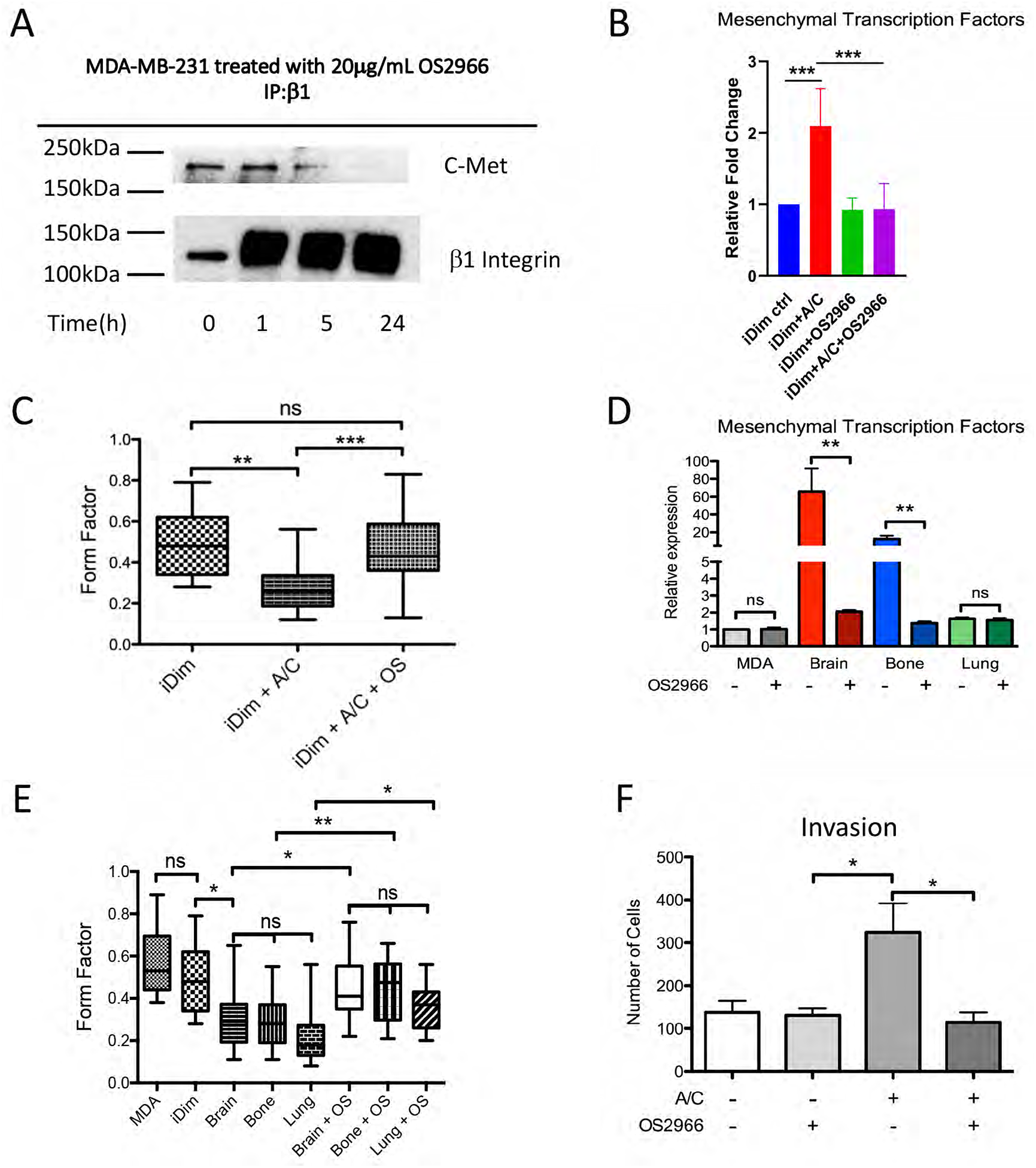
Pharmacologic disruption of c-Met/β1 binding reduced mesenchymal and stem cell profile of breast cancer cells. (**A**) Pharmacologic targeting of the c-Met/β1 complex with the OS2966 antibody disrupted complex formation as evidenced by IP. (**B**) AP21967 increased mesenchymal transcription factor expression in cultured MDA-MB-231-iDimerize-c-Met-β1 cells (P<0.001), changes reversed by OS2966 (n=3/group; P<0.001). (**C**) AP21967-induced c-Met/β1 complex formation lowered the form factor of MDA-MB-231-iDimerize-c-Met-β1 cells in culture (P=0.002), and this effect was reversed by OS2966 (P<0.001) (n=22/group). (**D**) MDA-MB-231-BO and MDA-MB-231-BR organ-seeking cells had higher expression of mesenchymal transcription factors TWIST, SNAIL, FOXC1, FOXC2, SLUG, and GSC than parental MDA-MB-231 cells, differences that were reversed with OS2966 treatment (n=3/group; P=0.008-0.009). (**E**) These organ-seeking cells also had lower form factor than parental MDA-MB-231 cells (P<0.001), differences that were reversed with OS2966 treatment (P=0.007-0.03) (n=22-24/group). (**F**) AP21967 increased invasiveness of MDA-MB-231-iDimerize-c-Met-β1 cells in culture (P=0.047), a change reversed by OS2966 (n=3/group; P=0.038). *P<0.05; **P<0.01; ***P<0.001.

## DISCUSSION

Although cancer death rates have declined over the past decade and survival have been prolonged in several cancer types with the development of novel therapeutics, patients with metastatic disease have not shared equally in these improvements (21). Despite intense efforts to understand the mechanisms underlying the metastatic cascade with the goal of uncovering effective therapeutic targets, minimal advances have been made in the treatment of metastatic cancer. For cancer patients, the majority of morbidity and mortality is associated with metastatic disease (22).

While c-Met and β1 integrin are each known to individually contribute to metastases (23, 24), the mechanisms through which these drive metastases or invasive resistance remain uncertain, as their high baseline expression levels do not change tremendously during acquisition of metastases (23, 24). We previously addressed this knowledge gap by identifying a structural complex between c-Met and β1 integrin formed at significantly higher levels in metastatic tumors relative to their primary tumors (7). Here, we build upon that observation by determining which steps of the metastatic cascade the c-Met/β1 integrin complex drives and whether the complex promotes organ-specific metastases.

As mentioned, the metastatic cascade comprises five major steps: local invasion at the primary site, intravasation, extravasation, invasive colonization of the metastatic site, and proliferation at the metastatic site (3). We previously demonstrated a role for the c-Met/β1 integrin complex in invasion at the primary site but not in proliferation at the metastatic site (7). Here, we build upon that finding by showing that the c-Met/β1 integrin complex promotes intravasation rather than extravasation, and enhances organ-specific invasive colonization of metastatic sites, and that these processes can not only be induced by activating c-Met/β1 integrin complex formation but can be reversed by genetic or pharmacologic targeting of the complex.

Our observation that the c-Met/β1 integrin complex drives intravasation into the circulation likely reflects our demonstration of the complex promoting breast cancer cell adhesion to endothelial cells. This finding is consistent with a demonstrated role of β1 integrin in tumor cell adhesion to endothelial cells (25) via binding of tumoral α4β1 integrin to endothelial VLA-4. Because the c-Met/β1 integrin complex promotes ligand-independent conformational changes in β1 integrin that structurally resemble activation (7) it is likely that the c-Met/β1 complex promotes β1 integrin functionality in general, including VLA-4 binding. Interestingly, our demonstration that the c-Met/β1 integrin complex does not promote extravasation suggests distinct mediators of tumor cell trafficking into versus out of circulation.

Following extravasation, cancer cells need to find metastatic sites, which tumor cells accomplish through a variety of mediators currently being characterized (6). Our study helped address this knowledge gap because we found that cancer cells with induced c-Met/β1 complex formation demonstrated organ specific metastasis with a preference for osseous colonization. This was evident from a significantly higher burden of osseous micrometastases after *in vivo* c-Met/β1 complex induction when organ specific qPCR was performed. This is likely due to preferential affinity of the c-Met/β1 complex for collagen type I, the primary ECM component of the bone. This finding of molecular alterations promoting organ-specific metastases is consistent with the observation that cancer cells target specific organs for metastases based on preferential affinity for organ-specific ECM (26). Even our understanding that cancer cells can secrete factors that can prime pre-metastatic niches in remote organs is based on the presumption that these tumor cell secreted factors bring immune cells into the pre-metastatic niche to remodel the ECM to make it more conducive to metastases (27). This finding makes the c-Met/β1 complex an even more appealing therapeutic target as bone is the third most frequent site of metastases, with osseous metastases conferring a poor prognosis (28).

Based on our findings, the promotion of increased metastases by the c-Met/β1 integrin complex is driven by the Wnt and hedgehog signaling pathways. These pathways have been implicated as mediators of metastases (29), although conflicting evidence exists as to whether they work synergistically or antagonistically in this process (30). Further work will be needed to delineate the sequence of events from c-Met/β1 integrin complex formation that precipitates Wnt and hedgehog pathway signaling and the associated downstream events we identified such as enrichment of stem cell fraction and mesenchymal transcription factor expression.

Our findings have significant translational implications in both preventing promotion of aggressive cancer behavior and preventing metastases. We found that bevacizumab increased c-Met/β1 integrin complex formation in breast cancer cells, a potential mechanism of the preclinical observation that VEGF-targeted therapies (31) and bevacizumab (32) increase the metastatic potential of cancer cells. Importantly, our study demonstrates that there are viable therapeutic agents to inhibit c-Met/β1 complex formation and its downstream events. Specifically, our demonstration that OS2966, a neutralizing antibody that disrupts the ability of β1 integrin to bind c-Met, offset the changes induced by c-Met/β1 integrin complex formation in cultured cells is an encouraging finding that warrants further evaluation.

## METHODS

### Study Approval

Human tissue research was approved by the UCSF IRB (approval #11-06160). Animal experiments were approved by the UCSF IACUC (approval #AN105170-02).

### Cell culture

MDA-MB-231 human breast adenocarcinoma cells (ATCC HTB-26) were passaged fewer than six months and verified by providing sources using short tandem repeat (STR) profiling and confirmed to be Mycoplasma free. MDA-MB-231-iDimerize-c-Met-β1 cells containing the Lenti-X iDimerize inducible heterodimer system were created as described previously (7). Complex formation was induced in these cells by treating with AP21967 (A/C ligand heterodimerizer; Takara Bio). MDA-MB-231-iDimerize-c-Met-β1/luc cells were created by lentiviral transduction of MDA-MB-231-iDimerize-c-Met-β1 cells and subsequent confirmation of bioluminescence. MDA-MB-231-BR, MDA-MB-231-BS, and MDA-MB-231-LS brain, bone-, and lung-seeking cells were provided by Dr. Joan Massagué (Memorial Sloan Kettering; New York, NY) (16–18). Some cells were treated with β1 neutralizing antibody OS2966 (kindly provided by Oncosynergy; Greenwich, CT) or VEGF neutralizing antibody bevacizumab (UCSF pharmacy). MDA-MB-231 cells were transduced to express CRISPR/Cas9 followed by transduction with guide RNAs targeting β1 integrin, with Western blot confirming β1 knockdown. The resulting MDA-MB-231/CRISPRβ1 cells were then transduced with lentiviral vectors expressing wild type β1 integrin or β1 integrin with change of amino acid 246 or 287 from aspartate to alanine (β1D246A), leading to creation of MDA-MB-231/wtβ1 and MDA-MB-231/β1D246A cells. Breast cancer cells were cultured in DMEM/F-12 supplemented with 10% fetal bovine serum and 1% penicillin/streptomycin and passaged for less than 6 months. Human umbilical vein endothelial cells (HUVECs) prescreened for angiogenesis were cultured in EGM-2 (Lonza). Breast cancer cells were cultured in MammoCult (Stemcell Technologies) on non-TC treated flasks to form mammospheres. Media collected from these cells was utilized as breast cancer stem cell conditioned media.

### Intravasation and extravasation transwell assays

#### Intravasation

In modification of a previously described protocol (33), 8-micron pore transwell inserts (Corning) were *inverted* and coated with 100uL of 6 μg/mL of growth factor reduced Matrigel (Corning) in DPBS (Gibco) for 1 hour at room temperature. Excess Matrigel was removed, and 1×10^6^ HUVECs stained with CellTracker Green CMFDA (Invitrogen) were plated on the Matrigel-coated inverted transwells in 100 μL of EGM-2 for 4 hours in a 37°C CO2 incubator to create a monolayer as previously described (33). Transwells were then flipped right side up into a 24 well plate (Corning) and 30,000 tumor cells stained with CellTracker Orange CMRA (1:1000) (Invitrogen) were added. Transwells were washed with PBS and fixed in 4% paraformaldehyde (PFA)/PBS for 15 minutes. Inserts were mounted on a slide with DAPI Fluoromount-G (Southern Biotech). Only tumor cells that breached the endothelial monolayer were scored as a positive transendothelial migration event. Nine fields of view were acquired for each transwell at a 20X magnification.

#### Extravasation

In modification of a previously described protocol (34), 8-micron pore FluoroBlok transwell inserts (Corning) were coated inside the chamber with 100 μL of 6 μg/mL of growth factor reduced Matrigel (Corning) in DPBS (Gibco) for 1 hour at room temperature. Excess Matrigel was removed, and 1×10^6^ HUVECs stained with CellTracker Green CMFDA (Invitrogen) were plated on the Matrigel-coated FluoroBlok transwells in 100μL of EGM-2 for 4 hours in a 37°C CO_2_ incubator. 30,000 tumor cells stained with CellTracker Orange CMRA (Invitrogen) were added. Transwells were washed with PBS and fixed in 4% PFA/PBS for 15 minutes. Inserts were mounted on a slide with DAPI Fluoromount-G (Southern Biotech). Only tumor cells that were on a different plane from the endothelial monolayer were scored as a positive extravasation event. Nine fields of view were acquired for each transwell at a 20X magnification (34).

### HUVEC adhesion assays

48 well plates were coated with Matrigel as described above. HUVECs were seeded at a density of 50,000 cells per well and grown to confluency. Tumor cells were stained with CellTracker Green CMFDA (Invitrogen). EGM-2 media was removed and DMEM/F-12 was added for the remainder of the assay. 25,000 tumor cells were added to each well and incubated to allow adhesion to the HUVEC monolayer. After 30 minutes, media was removed and wells were washed three times with DPBS to remove any unbound tumor cells. Wells were fixed in 4% PFA/PBS for 15 minutes. 3 fields of view were acquired for each well (35).

### qRT-PCR

RNA extraction was performed using either Quick-RNA Miniprep kit (Zymo Research) or RNeasy Mini kit (Qiagen). Complementary DNA (cDNA) was synthesized using the qScript cDNA SuperMix (Quantabio). Quantitative RT-PCR using SYBR Green (Quantabio) was performed via QuantStudio 3 system (Applied Biosystems). Primers are listed in **Supplemental Table 4**.

### Proximity Ligation Assays

The Duolink Detection System (Sigma) was used to assess c-Met/β1 complex formation as previously described (7).

### Western Blot

Human tissue samples and cell preparations were harvested in Complete RIPA Buffer - 1× radio immunoprecipitation buffer (10× RIPA; 9806, Cell Signaling) and one tablet each of PhoStop and Complete Mini (04906845001 and 04693124001, Roche). Insoluble materials were removed by centrifugation at 300 × *g* for 20 min at 4°C. Protein concentration was determined using the bicinchronic acid (BCA) assay (23225, Thermo Scientific). Samples were prepared with 10–30 μg of protein in RIPA buffer with 4× LDS loading buffer (LP0001, Life Technologies). Samples were electrophoresed on SDS/PAGE gels, transferred to PVDF membranes, and probed with primary antibodies overnight at 4°C. Membranes were detected using HRP-conjugated secondary antibodies and imaged using radiographic film or the Odyssey Fc imaging system (LI-COR).

### Immunoprecipitation

Samples were prepared in 500 μL Complete RIPA buffer containing 1000–1500 μg of protein. For β_1_ IP, 50 μL of protein A magnetic bead slurry (73778S, Cell Signaling) were aliquoted per sample and washed with 500 μL of Complete RIPA buffer. Beads were incubated with rabbit monoclonal anti-β_1_ antibody (1:20; ab52971, Abcam) in 200 μL of 0.1% Triton-X 100 PBS for 15 min at room temperature (RT). Beads were magnetically precipitated and resuspended with sample lysate for incubation on a rotator (4°C overnight). Antibody-bound beads were magnetically separated from the lysate supernatant and washed three times in 500 μL of Pierce™ IP Lysis Buffer (Thermo Scientific). Samples were eluted by precipitating beads on a magnetic rack and resuspending in 40 μL of 3× blue sample buffer 30× reducing agent: 1.25 M DTT (1:30; 7722s, Cell Signaling). Resulting samples were heated at 95°C (5 min), centrifuged at 300 × *g* at RT, and magnetically precipitated. 20 μL of the supernatant was used for each SDS/PAGE electrophoresis. Blots were probed with primary and secondary antibodies.

### Flow Cytometry

MDA-MB-231-iDimerize-c-Met-β1 cells were treated with either 0.5 mM A/C ligand or the equivalent dilution (1:1000) of 100% ethanol for 3 hours. The cells were then washed with PBS and dissociated with TrypLE (Thermo Fisher Scientific) and washed with PBS. 1 × 10^6^ cells were resuspended in 100uL of PBS + 10% FBS and stained with anti-CD44-APC (Biolegend) and anti-CD24-FITC (Biolegend) or isotype controls at 4C for 30 minutes. The samples were washed with PBS once and re-suspended in 1mL PBS + 10% FBS. Flow cytometry was performed via Sony SH800 and data was analyzed using FlowJo (Ashland, OR).

### Animal Work

Athymic mice (Jackson Laboratory) were anesthetized using isoflurane and buprenorphine injected intraperitoneally at a dose of 0.1 mg/kg. The injection area was prepared with Betadine solution. Intracardiac inoculation was performed by passing a 30 gauge × 0.5 inch needle into the left ventricle. Proper location was confirmed by presence of arterial blood pulsating into the needle. 10^5^ MDA-MB-231-iDimerize-c-Met-β1/luc cells suspended in 100 μl serum-free media were injected into the left ventricle. Serial whole body bioluminescence imaging was performed to track systemic metastases.

### Cell Morphology (Form Factor)

Cells were plated on 8-well Lab-Tek™ II Chamber Slides. 24 hours later, media was aspirated, and the slide was fixed in 4% PFA for 15 minutes. The slide was then washed in PBS for 5 minutes three times. The slide was then incubated in 6.6 μM Phalloidin (Cell Signaling Technology) diluted 1:20 in PBS for 15 minutes at RT. Subsequently the slide was washed in PBS once and mounted with DAPI Fluoromount-G (Southern Biotech). Slides were imaged at 20X magnification and form factor was analyzed using the Shape Descriptors plugin on Image J.

### Nanostring Multiplex Transcriptomic Analysis

Using the RNeasy Mini kit (Qiagen). RNA was extracted from: (1) MDA-MB-231-iDimerize-c-Met-β1 cells treated with and without AP21967 for 3 hours; and (2) MDA-MB-231/CRISPRβ1 cells transduced to express wild type β1 integrin or β1 integrin with change of amino acid 246 to alanine. A bioanalyzer was used to determine quantity and quality of the RNA sample. 100 ng of RNA was used for the Cancer Pathways Panel and 175ng of RNA was used for the Cancer Progression Panel. RNA from each sample was hybridized with the codeset for 18 hours. 30ul of the reaction was loaded into the nCounter cartridge and run on the nCounter SPRINT Profiler.

### Statistics

For comparing continuous variables, ANOVA/t-test (parametric) or Kruskal Wallis/Wilcoxon rank sum test (non-parametric) were used, with analysis on SPSS (IBM, v24.0). Cox proportional hazards was used to identify contributions of individual variables to patient survival. Error bars are standard deviations. The threshold for statistical significance was *P*<0.05.

## ACKNOWLEDGEMENTS

Work was supported by funding to M.K.A.’s lab from the NIH (1R01CA227136 and 2R01NS079697). D.L. was supported by a CTSI TL1 postdoctoral fellowship and the Neurosurgery Research Education Foundation (NREF). A.J. was an HHMI fellow and was supported by the NIH (1F31CA203372-01). A.C. and S.S.S. were HHMI fellows. J.S. was supported by the UCSF Yearlong Inquiry Program (YIP).

