## Supplemental Materials for "Role of c-Met/β1 integrin complex in the metastatic cascade"

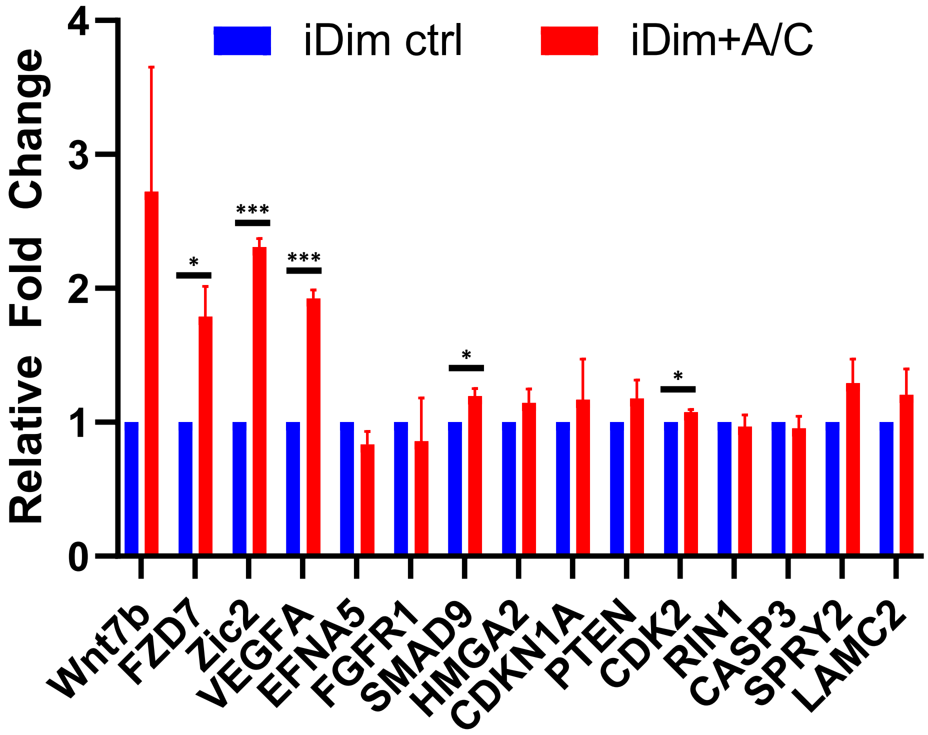


**Supplemental Figure 1. Validation of multiplex transcriptomic results by qPCR confirms elevated Wnt/hedgehog pathway signaling after c-Met/β1 complex formation. Related to Figures 1A-B.** qPCR was performed to validate results obtained from multiplex transcriptomic analysis. Bevacizumab increased expression of Fzd7 ($P=0.03$), Zic2 ($P<0.001$), VEGFA ($P<0.001$), Smad9 ($P=0.03$), and CDK2 ($P=0.03$). n=3/group. *$P<0.05$, **$P<0.01$, ***$P<0.001$.

**Supplemental Figure 2. Multiplex transcriptomic analysis reveals alteration in cancer pathways after c-Met/β1 complex formation. Related to Figure 1C.** Multiplex transcriptomic analysis revealed that treatment ot MDA-MB-231-iDimerize-c-Met-β1 cells with AP21967 (A/C ligand) upregulated several cancer pathways.

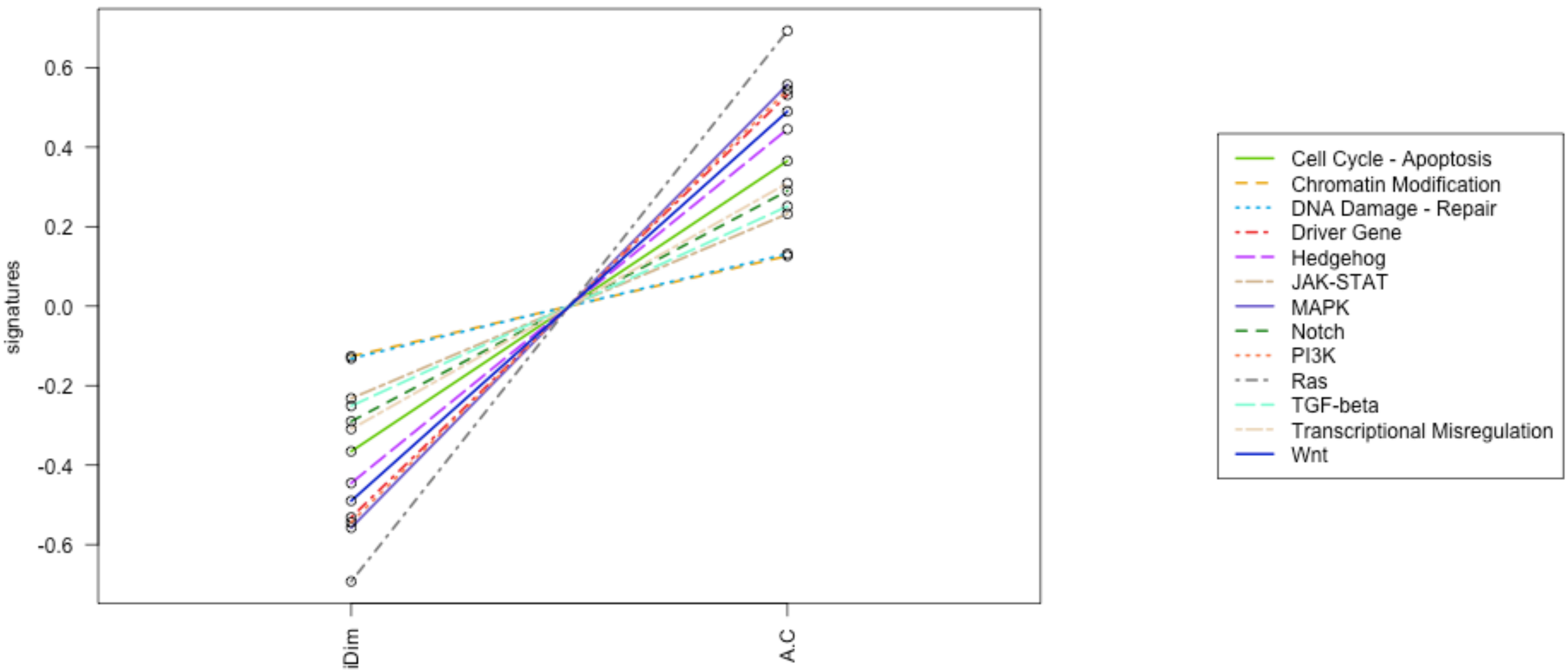

**Supplemental Figure 3. Individual mesenchymal transcription factors are upregulated after c-Met/β1 complex formation. Related to Figure 1D.** Shown are qPCR results for six mesenchymal transcription factors in MDA-MB-231-iDimerize-c-Met-β1 cells with and without 0.5 nM AP21967 (A/C ligand) for 24 hours. Results are integrated into a single cumulative mesenchymal transcription factor level in **Figure 1D** (P=0.013; n=3/group).

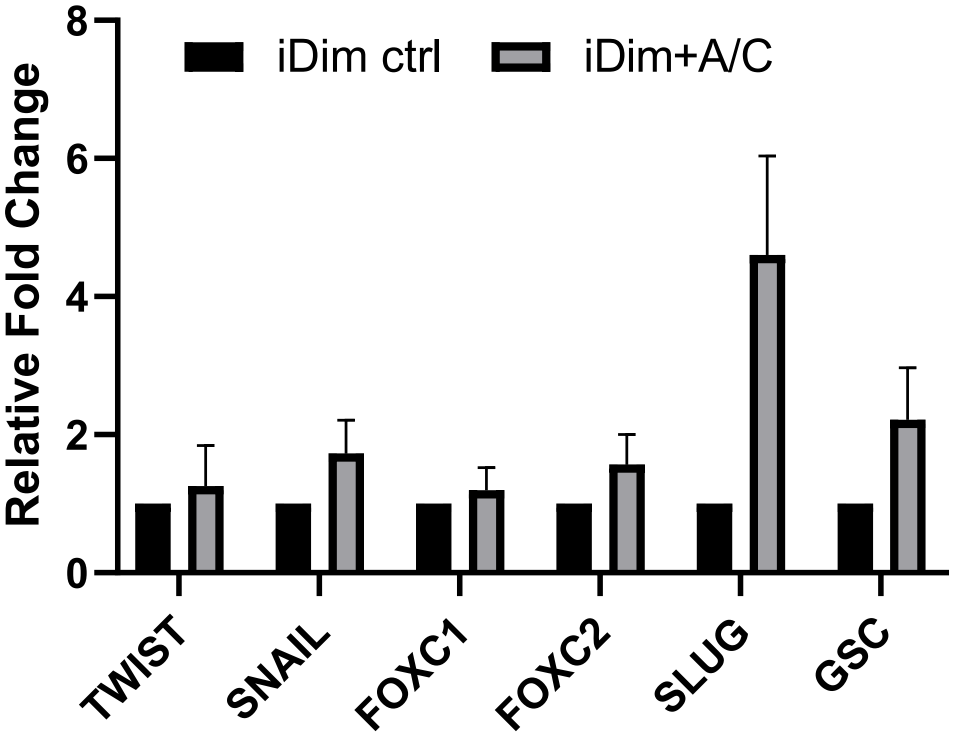

**Supplemental Figure 4. Pathways induced by c-Met/β1 complex formation. Related to Figure 1E.** In addition to the four pathways shown in **Figure 1E**, c-Met/β1 complex formation in MDA-MB-231-iDimerize-c-Met-β1 cells induced by 0.5 nM AP21967 (A/C ligand) for 24 hours increased transcription of factors in the 6 pathways shown below.

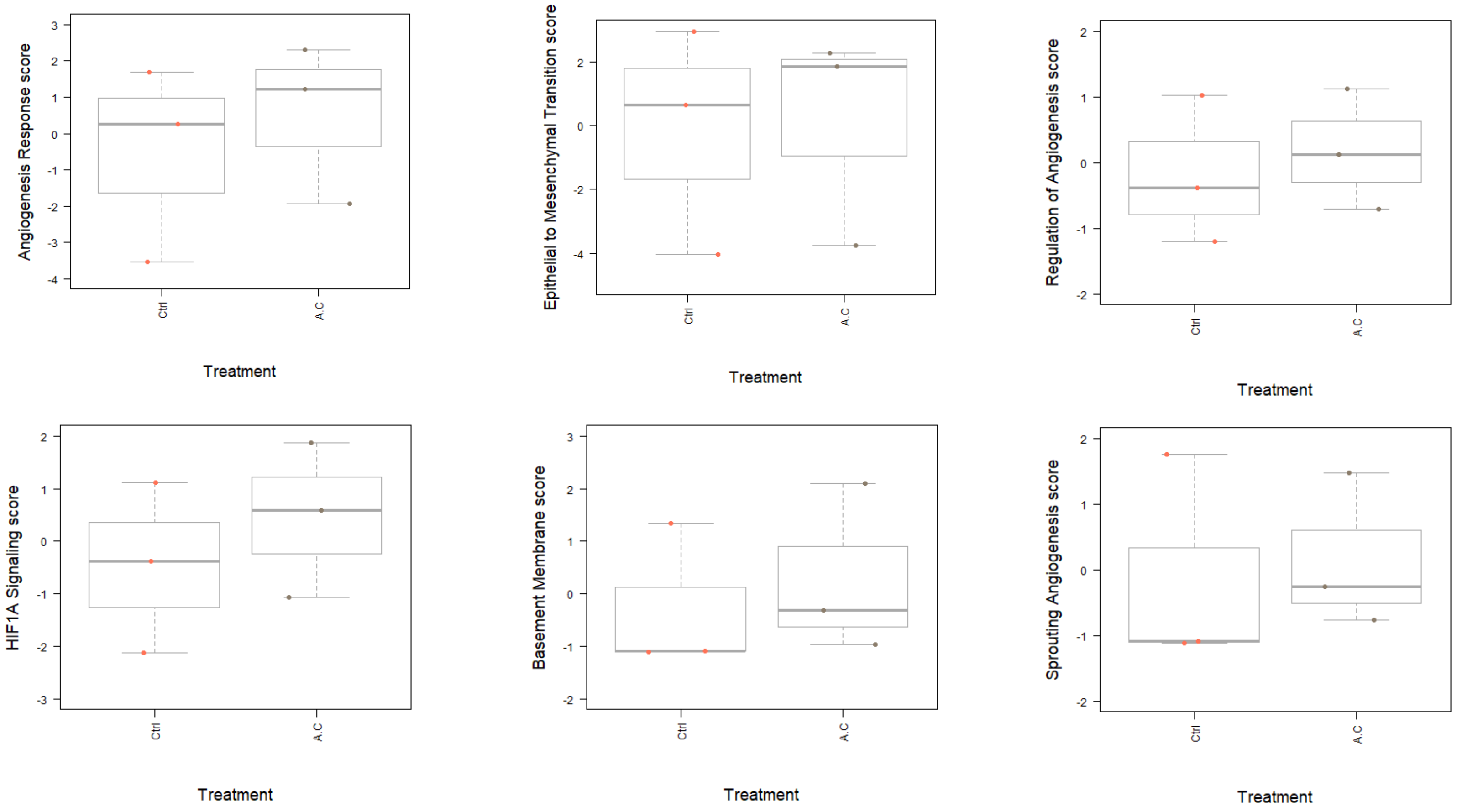

**Supplemental Figure 5. Individual stem cell genes upregulated after c-Met/β1 complex formation. Related to Figure 1F.** Shown are qPCR results for individual stem cell gene expression in MDA-MB-231-iDimerize-c-Met-β1 cells with and without 0.5 nM AP21967 (A/C ligand) for 3, 24, and 48 hours. The pooled aggregate fold change in expression of these stem cell genes at each time point is shown in **Figure 1F**. *P<0.05.

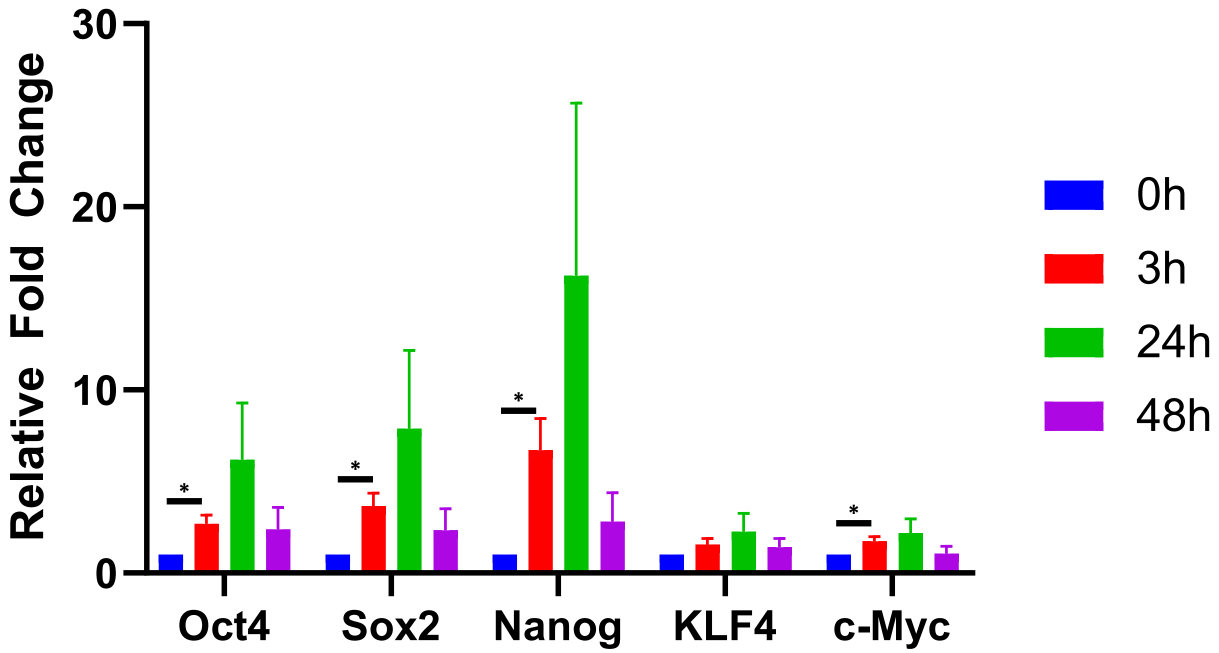

**Supplemental Figure 6. Confirming that mammospheres express breast cancer stem cell genes. Related to Figure 1G.** Use of qPCR to verify that mammospheres derived from MDA-MB-231 breast cancer cells expressed breast cancer stem cell genes at higher levels than adherent MDA-MB-231 cells. Shown to the left are pooled results for all 5 genes, and to the right are individual results from the five stem cell genes. *P<0.05; **P<0.01; ***P<0.001.

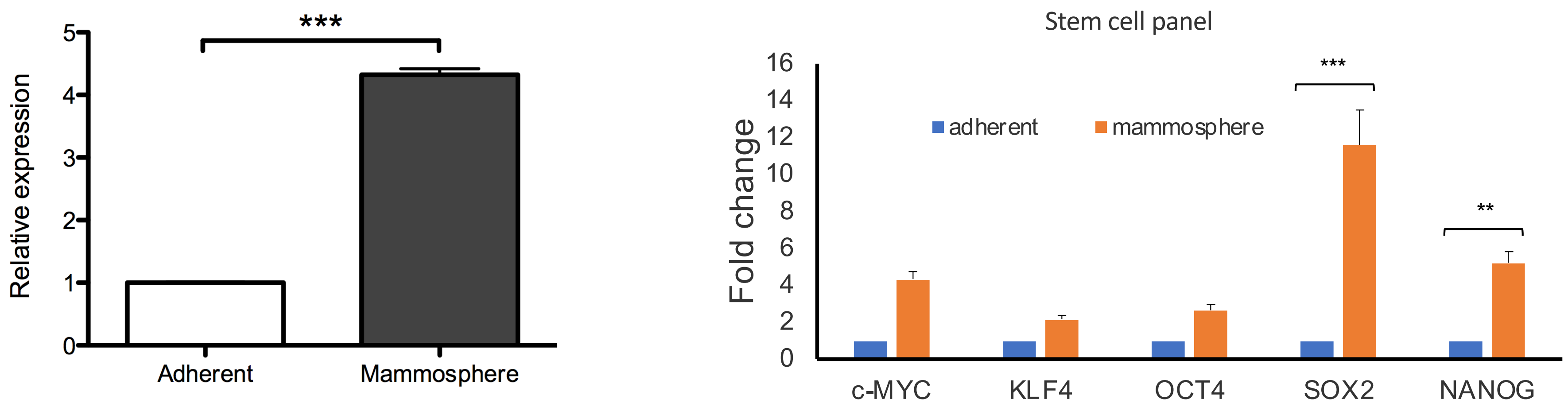

**Supplemental Figure 7. Mammosphere conditioned media promotes intravasation of breast cancer cells. Related to Figure 2D.** Shown are immunostainings of CMRA-labeled MDA-MB-231 breast cancer cells which were incubated in a cell culture intravasation assay for 48 hours in the absence or presence of mammosphere-conditioned media (MCM). Results are quantified in **Figure 2D**. n=3/group. Scale bar, 200 μm.

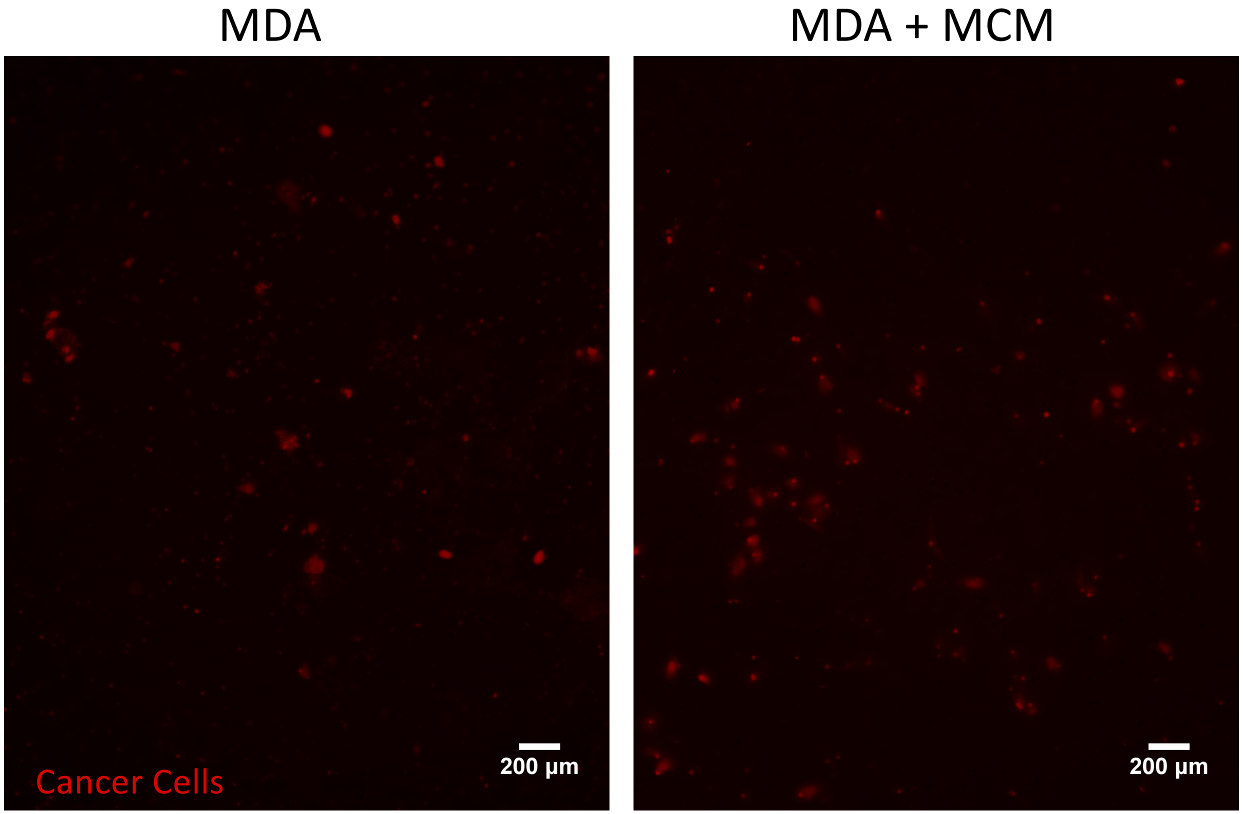

**Supplemental Figure 8. Bevacizumab increases c-Met/β1 complex formation in breast cancer cells.** Immunoprecipitation to pull down β1 integrin revealed more c-Met/β1 complex formation after treating MDA-MB-231 breast cancer cells with 2.5 μg/mL bevacizumab for 24 hours.

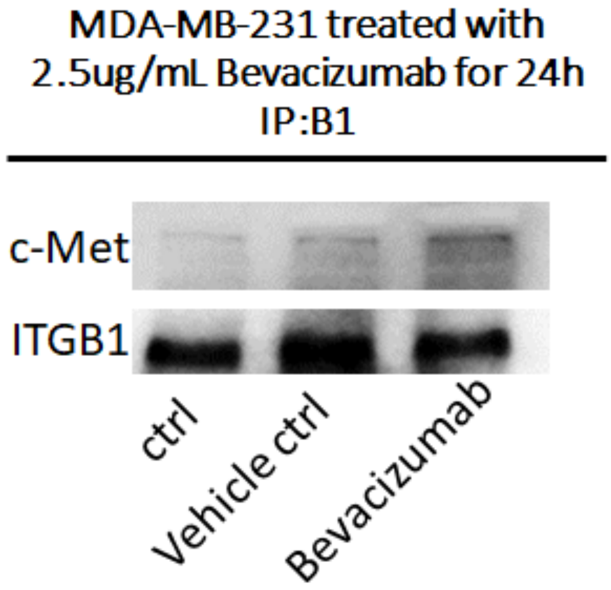

**Supplemental Figure 9. Bevacizumab increases intravasation of breast cancer cells. Related to Figure 2F.** Shown are immunostainings of CMRA-labeled MDA-MB-231 breast cancer cells and CMFDA-labeled HUVEC cells with DAPI nuclear staining in blue at the 48 hour time point after intravasation assays. n=3/group. Results are quantified in **Figure 2F**. Scale bar, 20 μm.

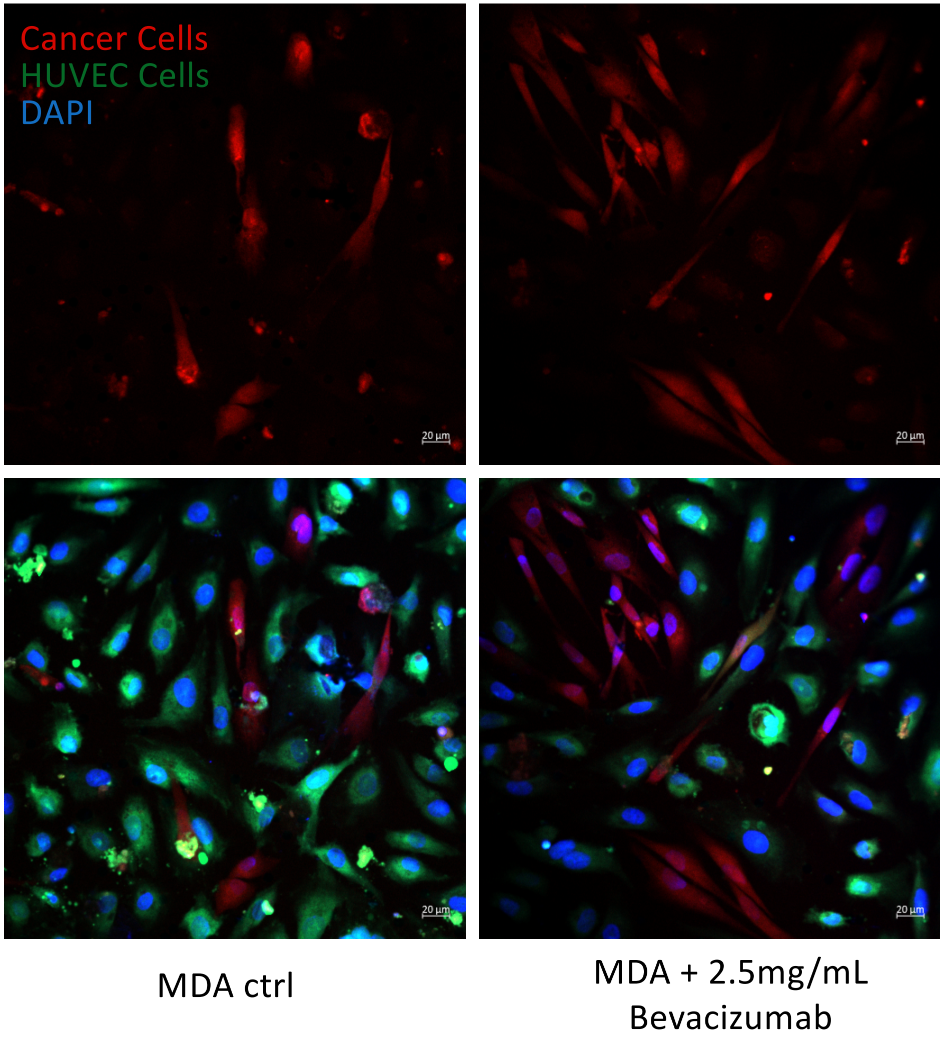

**Supplemental Figure 10. c-Met/β1 complex does not alter extravasation of breast cancer cells.** Shown are immunostainings of CMRA-labeled MDA-MB-231 cells at the 48 hour time point after a cell culture extravasation assay. Results are quantified in **Figure 2G**.

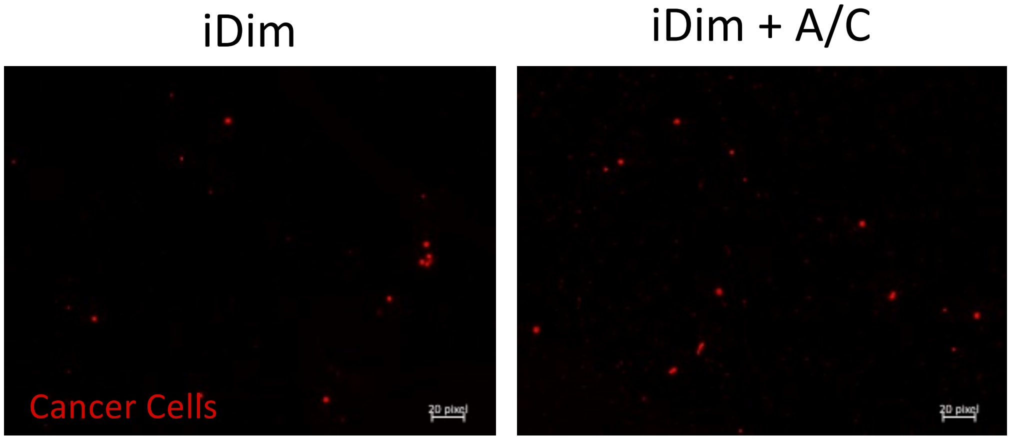

**Supplemental Figure 11. Proximity-ligation assays of patient samples reveal c-Met/β1 complexes in different types of metastases**. Related to **Figure 3G.** Shown are PLA immunostainings of breast, renal cell cancer (RCC), and prostate cancer metastases to brain versus bony structures. Results were quantified in **Figure 3G**.

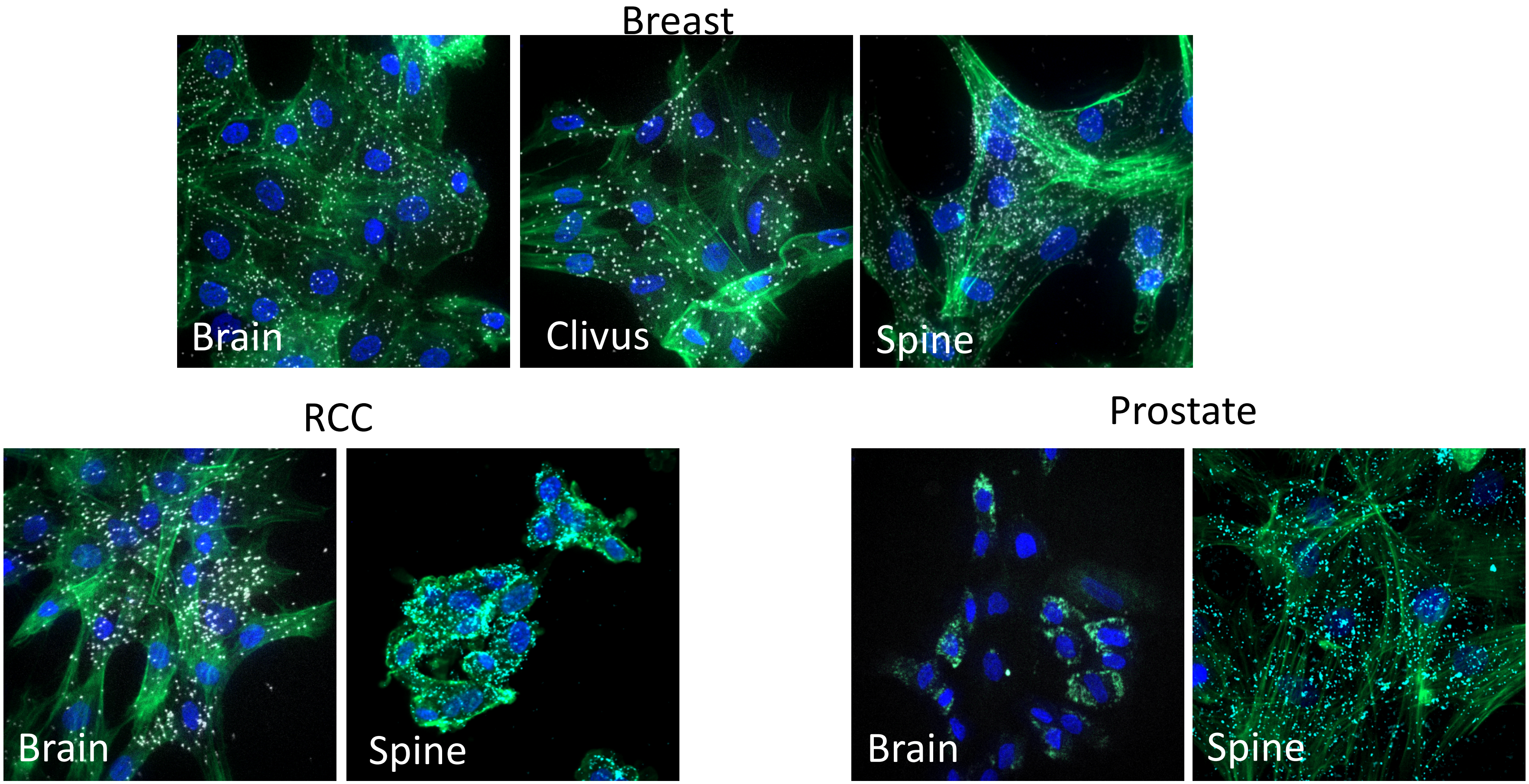

**Supplemental Figure 12. Immunoprecipitation of patient samples reveals c-Met/β1 complexes in different types of metastases.** Shown are immunoprecipitations of human tumors in which c-Met is precipitated and blotted for β1 integrin. The ratio of β1 integrin to c-Met quantified band intensity was higher in patient bony metastases (n=11) relative to brain metastases (n=12) from different patients (left two bars; $P<0.01$) and in paired bone metastases relative to brain metastases from the same patients (n=3; right panel; $P<0.001$). *$P<0.05$; **$P<0.01$; ***$P<0.001$.

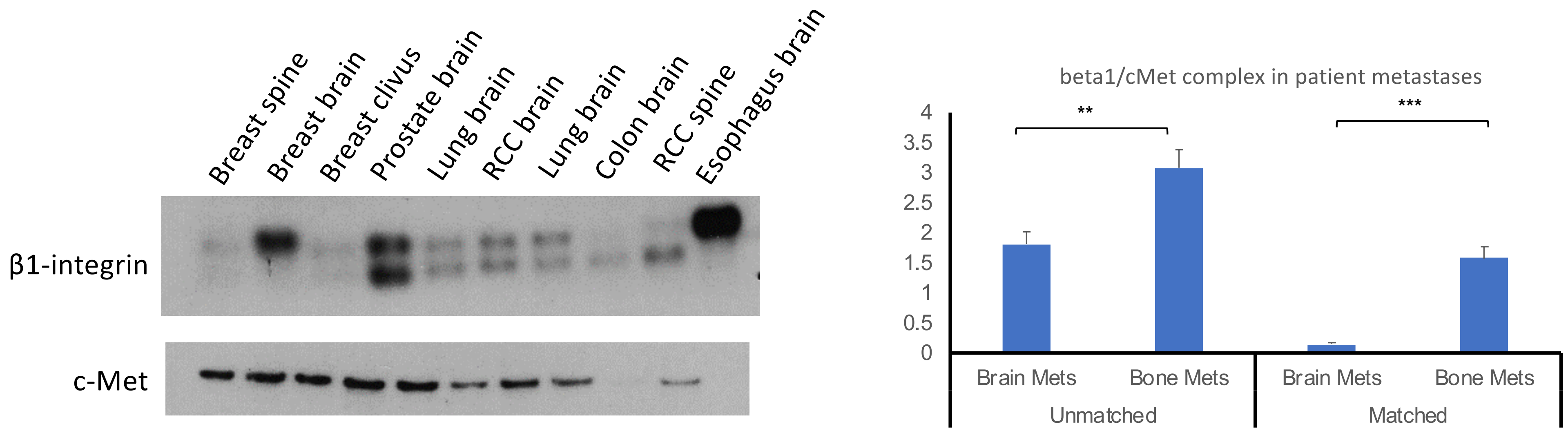

**Supplemental Figure 13. $\beta$1 integrin knockdown via CRISPRi in breast cancer cells.** MDA-MB-231 breast cancer cells were engineered to express KRAB CAS, followed by guide RNAs targeting $\beta$1 integrin. Western blot revealed loss of $\beta$1 integrin expression in the resulting cells.

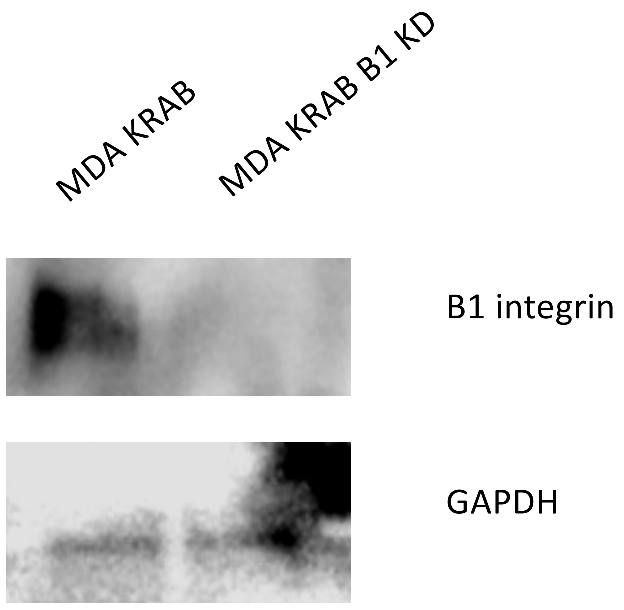

**Supplemental Figure 14. Immunoprecipitation reveals point mutations that lower binding of β1 integrin to c-Met.** Shown are results when immunoprecipitating for β1 integrin and blotting for c-Met as well as confirmatory blot for β1 integrin in MDA-MB-231 cells engineered for β1 integrin loss via CRISPR followed by restoration of wild-type β1 integrin (ctrl) or restoring β1 integrin with point mutations D246A and D287A which we previously demonstrated to reduce binding to c-Met in glioblastoma cells. Results confirmed reduced binding to c-Met in MDA-MB-231 breast cancer cells.

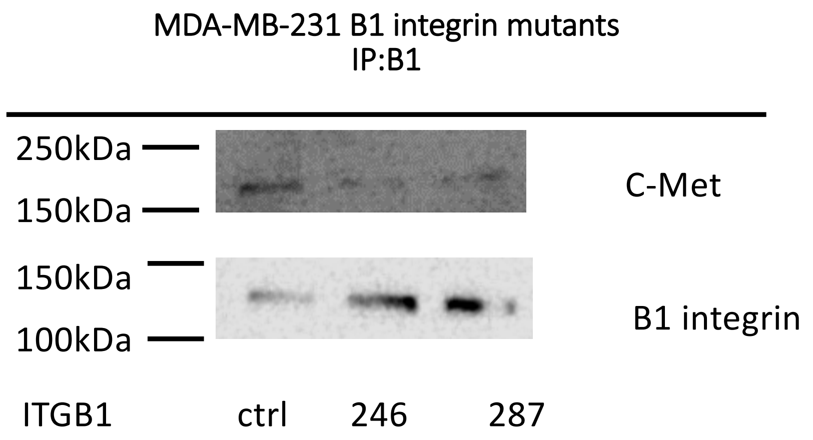

**Supplemental Figure 15. Individual mesenchymal transcription factors upregulated after c-Met/β1 complex formation are not upregulated when cells are treated with OS2966 blocking antibody.** Shown are qPCR results from MDA-MB-231-iDimerize-c-Met-β1 cells treated with 0.5 nM AP21967 (A/C ligand) for 3 hours and/or 20 μg/mL OS2966 for 24 hours. Aggregate results from these PCRs are presented in **Figure 5C**. *P<0.05.

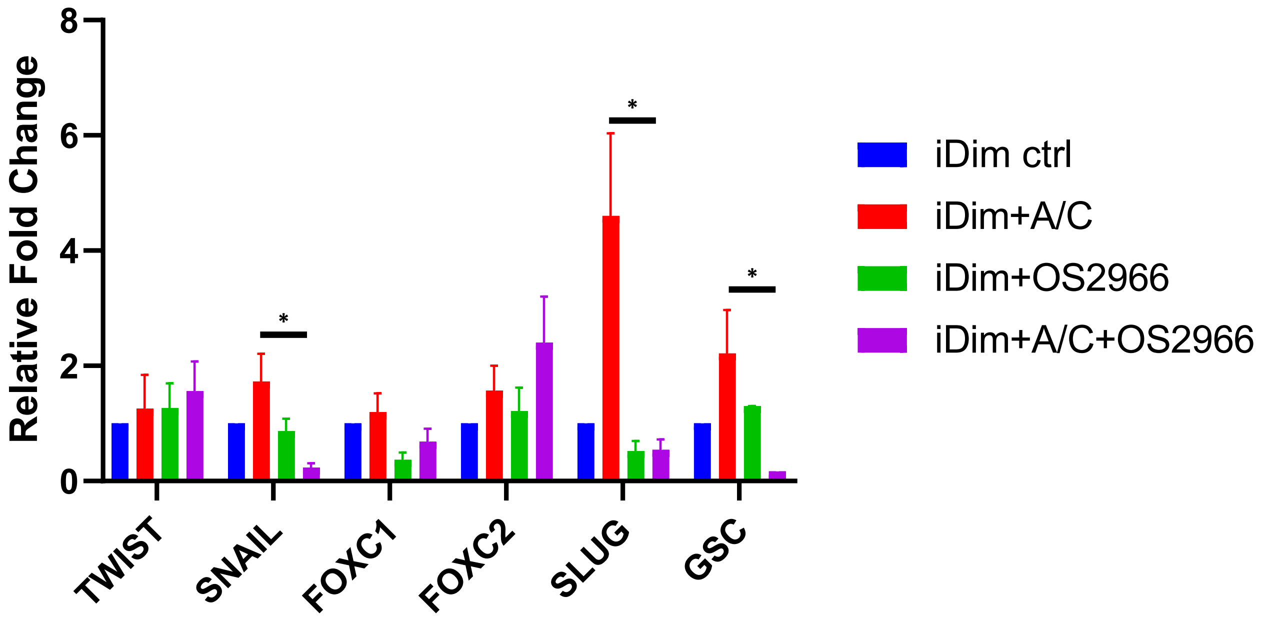

| | Log2 fold cha | std error (log | Lower confid | Upper confid | Linear fold ch | Lower confid | Upper confid | P-value | BY.p.value | method | probe.ID |
|---|---|---|---|---|---|---|---|---|---|---|---|
| ZIC2-mRNA | -0.38 | 0.0503 | -0.479 | -0.282 | 0.768 | 0.718 | 0.823 | 0.00164 | 1 | loglinear | NM_007129.2:1849 |
| CASP3-mRNA | -0.13 | 0.0196 | -0.168 | -0.0913 | 0.914 | 0.89 | 0.939 | 0.00272 | 1 | loglinear | NM_032991.2:685 |
| VEGFA-mRNA | -0.636 | 0.122 | -0.876 | -0.397 | 0.643 | 0.545 | 0.76 | 0.00649 | 1 | lm.nb | NM_001025366.1:1325 |
| CDK2-mRNA | 0.1 | 0.0205 | 0.0598 | 0.14 | 1.07 | 1.04 | 1.1 | 0.0082 | 1 | loglinear | NM_001798.2:220 |
| WNT7B-mRN | -0.744 | 0.14 | -1.02 | -0.471 | 0.597 | 0.494 | 0.722 | 0.0129 | 1 | Wald | NM_058238.1:1535 |
| FZD7-mRNA | -0.422 | 0.103 | -0.624 | -0.221 | 0.746 | 0.649 | 0.858 | 0.0148 | 1 | loglinear | NM_003507.1:1890 |
| CDKN1A-mRI | -0.587 | 0.148 | -0.877 | -0.298 | 0.666 | 0.545 | 0.813 | 0.0165 | 1 | lm.nb | NM_000389.2:1975 |
| SPRY2-mRNA | -0.334 | 0.0883 | -0.507 | -0.16 | 0.794 | 0.704 | 0.895 | 0.0195 | 1 | loglinear | NM_005842.2:85 |
| EFNA5-mRNA | -0.367 | 0.0993 | -0.562 | -0.172 | 0.775 | 0.677 | 0.887 | 0.0209 | 1 | loglinear | NM_001962.2:5035 |
| HMGA2-mRN | -0.342 | 0.0999 | -0.538 | -0.147 | 0.789 | 0.689 | 0.903 | 0.0266 | 1 | loglinear | NM_003484.1:328 |
| RIN1-mRNA | -0.218 | 0.0651 | -0.346 | -0.0905 | 0.86 | 0.787 | 0.939 | 0.0285 | 1 | loglinear | NM_004292.2:2572 |
| PTEN-mRNA | -0.247 | 0.0776 | -0.399 | -0.0952 | 0.842 | 0.758 | 0.936 | 0.0333 | 1 | loglinear | NM_000314.3:1675 |
| SMAD9-mRN | -0.44 | 0.124 | -0.683 | -0.196 | 0.737 | 0.623 | 0.873 | 0.0386 | 1 | Wald | NM_005905.2:1595 |
| LAMC2-mRN | -0.496 | 0.166 | -0.822 | -0.171 | 0.709 | 0.566 | 0.889 | 0.0405 | 1 | loglinear | NM_005562.2:2819 |
| FGFR1-mRNA | -0.389 | 0.133 | -0.649 | -0.129 | 0.764 | 0.638 | 0.915 | 0.0428 | 1 | loglinear | NM_015850.2:1335 |
| LTBP1-mRNA | -0.448 | 0.156 | -0.755 | -0.142 | 0.733 | 0.593 | 0.906 | 0.0456 | 1 | lm.nb | NM_000627.3:4124 |
| NBN-mRNA | -0.144 | 0.0503 | -0.242 | -0.0453 | 0.905 | 0.845 | 0.969 | 0.0459 | 1 | loglinear | NM_001024688.1:1105 |

**Supplemental Table S1. Genes related to canonical cancer pathways that are upregulated by c-Metβ1 compelx formation.** Shown are genes whose expression is changed when MDA-MB-231-iDimerize-c-Met-β1 cells were treated with AP21967 based on assessed in the NanoString nCounter platform using a 770 gene multiplex related to 13 cancer-associated canonical pathways

| | Undirected T | Directed Treatment: differential expression in AC vs. baseline of Ctrl |
|---|---|---|
| Cell Cycle - A | 1.293 | 0.949 |
| Chromatin M | 1.14 | 0.988 |
| DNA Damage | 0.901 | 0.432 |
| Driver Gene | 1.072 | 0.862 |
| Hedgehog | 2.742 | 2.742 |
| JAK-STAT | 1.121 | 1.115 |
| MAPK | 1.224 | 1.203 |
| Notch | 1.151 | 1.151 |
| PI3K | 1.376 | 1.095 |
| Ras | 1.245 | 1.225 |
| TGF-beta | 1.218 | 1.139 |
| Transcription | 1.153 | 0.966 |
| Wnt | 1.474 | 1.432 |

**Supplemental Table S2. Pathways activated by c-Met/β1 complex induction in breast cancer cells.** Shown are the pathways activated when MDA-MB-231-iDimerize-c-Met-β1 cells were treated with AP21967 based on assessed in the NanoString nCounter platform using a 770 gene multiplex related to 13 cancer-associated canonical pathways

| | Log2 fold cha | std error (log | Lower confid | Upper confid | Linear fold ch | Lower confid | Upper confid | P-value | BY.p.value | method | Gene.sets | probe.ID |
|---|---|---|---|---|---|---|---|---|---|---|---|---|
| PGK1-mRNA | 0.296 | 0.0663 | 0.166 | 0.426 | 1.23 | 1.12 | 1.34 | 0.0111 | 1 | loglinear | HIF1A Signali | NM_000291.2:1030 |
| AKAP2-mRNA | -0.246 | 0.0566 | -0.357 | -0.135 | 0.843 | 0.781 | 0.911 | 0.0122 | 1 | loglinear | Epithelial to | NM_001004065.4:4956 |
| FBN1-mRNA | -0.38 | 0.0877 | -0.552 | -0.208 | 0.768 | 0.682 | 0.866 | 0.0123 | 1 | loglinear | Basement M | NM_000138.3:6420 |
| ROCK1-mRNA | -0.141 | 0.0331 | -0.205 | -0.0758 | 0.907 | 0.867 | 0.949 | 0.0132 | 1 | loglinear | Cell Adhesion | NM_005406.1:2660 |
| TGFBR2-mRN | 0.194 | 0.0483 | 0.0997 | 0.289 | 1.14 | 1.07 | 1.22 | 0.0158 | 1 | loglinear | Cell Prolifera | NM_001024847.1:1760 |
| BMPR2-mRN | -0.215 | 0.0563 | -0.325 | -0.105 | 0.862 | 0.798 | 0.93 | 0.0188 | 1 | loglinear | Cell Prolifera | NM_001204.5:1875 |
| ARAP2-mRNA | -0.212 | 0.0557 | -0.321 | -0.103 | 0.863 | 0.8 | 0.931 | 0.019 | 1 | loglinear | Epithelial to | NM_015230.2:4875 |
| NAP1L3-mRN | 0.855 | 0.19 | 0.482 | 1.23 | 1.81 | 1.4 | 2.34 | 0.0205 | 1 | Wald | Epithelial to | NM_004538.4:1070 |
| VAMP8-mRN | 0.421 | 0.121 | 0.183 | 0.659 | 1.34 | 1.14 | 1.58 | 0.0257 | 1 | loglinear | Epithelial to | NM_003761.3:260 |
| AGR2-mRNA | 1.46 | 0.424 | 0.625 | 2.29 | 2.74 | 1.54 | 4.88 | 0.0264 | 1 | lm.nb | Epithelial to | NM_006408.3:580 |
| ENO2-mRNA | 0.228 | 0.067 | 0.0962 | 0.359 | 1.17 | 1.07 | 1.28 | 0.0274 | 1 | loglinear | HIF1A Signali | NM_001975.2:1855 |
| COL7A1-mRN | -0.516 | 0.134 | -0.779 | -0.253 | 0.7 | 0.583 | 0.839 | 0.0311 | 1 | Wald | Basement M | NM_000094.2:390 |
| CDH11-mRNA | 0.386 | 0.119 | 0.153 | 0.62 | 1.31 | 1.11 | 1.54 | 0.0315 | 1 | loglinear | Cell Adhesion | NM_001797.2:1835 |
| CXCR4-mRNA | -0.214 | 0.0665 | -0.344 | -0.084 | 0.862 | 0.788 | 0.943 | 0.0321 | 1 | loglinear | Epithelial to | NM_003467.2:1335 |
| EPAS1-mRNA | 0.474 | 0.158 | 0.165 | 0.783 | 1.39 | 1.12 | 1.72 | 0.0397 | 1 | lm.nb | Angiogenesis | NM_001430.3:4246 |
| RPS6KB2-mR | 0.186 | 0.0624 | 0.0634 | 0.308 | 1.14 | 1.04 | 1.24 | 0.0409 | 1 | loglinear | Cellular Grov | NM_003952.2:980 |
| MED1-mRNA | -0.154 | 0.0544 | -0.26 | -0.0469 | 0.899 | 0.835 | 0.968 | 0.0477 | 1 | loglinear | Angiogenesis | NM_004774.3:806 |

**Supplemental Table S3. Genes whose expression is altered when β1 integrin cannot bind c-Met in breast cancer cells.** Shown are the genes whose expression is altered when MDA-MB-231 breast cancer cells undergo CRISPRi knockdown of β1 integrin followed by lentiviral transduction of the β1D246A mutant vs. wild-type β1 integrin, as assessed in the NanoString nCounter platform using a 770 gene multiplex related to each step in the cancer progression process.

**Supplementary Table S4. Primers used for qPCR.** Shown are primers used for qPCR to assess expression of genes in breast cancer cells.

| Gene Target | Forward | Reverse |
|---|---|---|
| STEM CELL PANEL | | |
| *c-Myc* | *5'*- CAT CGT AAA CAC CAA CGT GC-3' | 5'- CCG CGT TCA TGT CGT AAT AG- 3' |
| *Klf4* | 5' –CAC CAT GCC GAT GTT CAT CGT AAA - 3' | 5' –TTA GGC GAA GGT GGA GTT GT - 3' |
| *Oct4* | 5' – CTT GCC TTG CTG CTC TAC CT – 3' | 5' – CAC ACA GGA TGG CTT GAA GA – 3' |
| *Sox2* | 5' – CAG CCA GAT GCA ATC AAT GC-3' | 5-GCA CTG AGA TCT TCC TAT TGG TGA A-3' |
| *Nanog* | 5'- GAC AAG CCA CAA GCT GAA CA-3' | 5'- GAG CCC ACA ATG GGA GAGT A-3' |
| NANOSTRING VALIDATION | | |
| *Wnt7B* | 5'-AGC CAA CAT CAT CTG CAA CA-3' | 5'-CTG GTA CTG GCA CTC GTT GA-3' |
| *Fzd7* | 5'-CGC CTC TGT TCG TCT ACC TC-3' | 5'-CCA TGA GCT TCT CCA GCT TC-3' |
| *Zic2* | 5'-AAT CCC AAG AAG AGC TGC AA-3' | 5'-ACA CTC CTC CCA GAA GCA GA-3' |
| *VEGFA* | 5'-AGG CCA GCA CAT AGG AGA GA-3' | 5'-TTT CTT GCG CTT TCG TTT TT-3' |
| *CDKN1A* | 5'-GAC ACC ACT GGA GGG TGA CT-3' | 5'-CCA CAT GGT CTT CCT CTG CT-3' |
| *CDK2* | 5'-TTG TCA AGC TGC TGG ATG TC-3' | 5'-TGA TGA GGG GAA GAG GAA TG-3' |
| *CASP3* | 5'-TTT TTC AGA GGG GAT CGT TG-3' | 5'-CGG CCT CCA CTG GTA TTT TA-3' |
| *LAMC2* | 5'-GGC TGG TCT TAC TGG AGC AG-3' | 5'-CAT CAG CCA GAA TCC CAT CT-3' |
| *SMAD9* | 5'-CCA CAG AAG CCT CTG AGA CC-3' | 5'-CCC AAC TCG GTT GTT CAG TT-3' |
| *EFNA5* | 5'-ATG TGT GTG TTC AGC CAG GA-3' | 5'-GGG CAG AAA ACA TCC AGG TA-3' |
| *FGFR1* | 5'-CGA TGT GCA GAG CAT CAA CT-3' | 5'-TGC TGG TTA CGC AAG CAT AG-3' |
| *HMGA2* | 5'-CCT AAG AGA CCC AGG GGA AG-3' | 5'-AAC TTG TTG TGG CCA TTT CC-3' |
| *PTEN* | 5'-CGA CGG GAA GAC AAG TTC AT-3' | 5'-AGG TTT CCT CTG GTC CTG GT-3' |
| *RIN1* | 5'-CCC AGA CCT AGT CCA GCT CA-3' | 5'-GGA GCT CCA GAA CTC AAT GC-3' |
| *SPRY2* | 5'-ATC AGA GCC ATC CGA AAC AC-3' | 5'-CAG ACC GTG GAG TCT CTC GT-3' |

| | | |
|---|---|---|
| MESENCHYMAL TRANSCRIPTION FACTOR PANEL | | |
| *GSC* | 5'-TCT CAA CCA GCT GCA CTG TC | 5'-GGC GGT TCT TAA ACC AGA CC-3' |
| *FOXC1* | 5'-CAT CCG CCA CAA CCT CTC GCT-3' | 5'-GTG CAG CCT GTC CTT CTC CTC C-3' |
| *FOXC2* | 5'-GCC TAA GGA CCT GGT GAA GC-3' | 5'-TTG ACG AAG CAC TCG TTG AG-3' |
| *ZEB1* | 5'-GCA CCT GAA GAG GAC CAG AG-5' | 5'-TGC ATC TGG TGT TCC ATT TT-3' |
| *ZEB2* | 5' GAC AGA TCA GCA CCA AAT GC-3' | 5'-GCT GAT GTG CGA ACT GTA GG-3' |
| *SLUG* | 5'-CAC TAT GCC GCG CTC TTC-3' | 5'-GGT CGT AGG GCT GCT GGA A-3' |
| *SNAIL* | 5'-TGG TTG CTT CAA GGA CAC AT-3' | 5'-GTT GCA GTG AGG GCA AGA A-3' |
| *TWIST* | 5'-GGA GTC CGC AGT CTT ACG AG-3' | 5'-TCT GGA GGA CCT GGT AGA GG-3' |
| ANALYZING TISSUES FOR MICROMETASTASES | | |
| *Luciferase* | 5’-GTG GTG TGC AGC GAG AAT AG-3’ | 5’-CGC TCG TTG TAG ATG TCG TTA G -3’ |
| STANDARDS | | |
| *GAPDH* | 5'-CAA TGA CCC CTT CAT TGA CC-3' | 5'-TTG ATT TTG GAG GGA TCT CG-3' |
| *ACTB* | 5'-GAG CAC AGA GCC TCG CCT TT-3' | 3'-ACA TGC CGG AGC CGT TGT C-3' |
